# The SARS-CoV-2 protein ORF3c is a mitochondrial modulator of innate immunity

**DOI:** 10.1101/2022.11.15.516323

**Authors:** Hazel Stewart, Yongxu Lu, Sarah O’Keefe, Anusha Valpadashi, Luis Daniel Cruz-Zaragoza, Hendrik A. Michel, Samantha K. Nguyen, George W. Carnell, Nina Lukhovitskaya, Rachel Milligan, Irwin Jungreis, Valeria Lulla, Andrew D. Davidson, David A. Matthews, Stephen High, Peter Rehling, Edward Emmott, Jonathan L. Heeney, James R. Edgar, Geoffrey L. Smith, Andrew E. Firth

## Abstract

The SARS-CoV-2 genome encodes a multitude of accessory proteins. Using comparative genomic approaches, an additional accessory protein, ORF3c, has been predicted to be encoded within the ORF3a sgmRNA. Expression of ORF3c during infection has been confirmed independently by ribosome profiling. Despite ORF3c also being present in the 2002-2003 SARS-CoV, its function has remained unexplored. Here we show that ORF3c localises to mitochondria during infection, where it inhibits innate immunity by restricting IFN-β production, but not NF-κB activation or JAK-STAT signalling downstream of type I IFN stimulation. We find that ORF3c acts after stimulation with cytoplasmic RNA helicases RIG-I or MDA5 or adaptor protein MAVS, but not after TRIF, TBK1 or phospho-IRF3 stimulation. ORF3c co-immunoprecipitates with the antiviral proteins MAVS and PGAM5 and induces MAVS cleavage by caspase-3. Together, these data provide insight into an uncharacterised mechanism of innate immune evasion by this important human pathogen.

## INTRODUCTION

Severe acute respiratory syndrome coronavirus 2 (SARS-CoV-2) is the causative agent of COVID-19 and the recent pandemic that has had unprecedented effects upon humanity, ranging from numerous casualties to severe economic impact. It is imperative to have a thorough understanding of the pathogen-host interactions that occur during infection. A key first step is to delineate and characterise functionally the full complement of accessory proteins encoded in the SARS-CoV-2 genome. However, most studies have been informed by pre-existing research based upon the closely related SARS-CoV (referred to herein as SARS-CoV-1 to avoid confusion), which caused a relatively minor outbreak in 2002 – 2003. As a result, proteins that had not been annotated in studies of SARS-CoV-1 were overlooked during the initial scientific response to the COVID-19 pandemic.

SARS-CoV-2 and SARS-CoV-1 belong to the taxon *Sarbecovirus*, a subgenus of the genus *Betacoronavirus* in the family *Coronaviridae*. Members of the *Coronaviridae* possess large positive-sense, single-stranded RNA genomes (approximately 30 kb in size) which contain 7 - 16 protein-coding open reading frames (ORFs). The two largest ORFs, ORF1a and ORF1b, are translated from the genomic mRNA directly (with translation of ORF1b depending on ribosomes making a programmed frameshift near the end of ORF1a) and encode polyproteins pp1a and pp1ab, which are cleaved into individual functional proteins that support viral replication. The remainder of the ORFs are translated from a set of “nested” subgenomic mRNAs (sgmRNAs). The encoded proteins are either structural components of the virion or so-called “accessory” proteins: mostly dispensable for viral replication in cell culture, these latter proteins nonetheless confer significant advantages to replication in hosts, often due to interactions with innate immune pathways. The assemblage of accessory proteins varies substantially across the *Coronaviridae*, and can vary even between closely related coronaviral species, thus constituting an important field of research. The potential translational repertoire of SARS-CoV-2 has been the subject of many publications, with multiple groups reporting evidence of novel ORFs via a range of approaches^1-8^. Collectively these studies highlight the lack of an established, accepted SARS-CoV-2 viral proteome because the functional relevance of these reports is seldom validated experimentally^9^.

In 2020, making use of the ∼54 then-available sarbecovirus genomes, we and others used comparative genomic approaches to analyse the coding capacity of SARS-CoV-2. Our analysis revealed a previously undetected conserved ORF, overlapping ORF3a in the +1 reading frame, and precisely coinciding with a region of statistically significantly enhanced synonymous site conservation in ORF3a-frame codons, indicative of a functional overlapping gene, ORF3c^10^. ORF3c was predicted independently by Cagliani et al. who termed it ORF3h^11^ and Jungreis et al. who termed it ORF3c^2^. A community consensus fixed the name as ORF3c^12^. Given ORF3c was previously unknown, it had not been the focus of any pre-existing studies despite being present in SARS-CoV-1. However its continued presence throughout evolution indicates that it is beneficial to successful viral replication, immune evasion or transmission, at least in natural hosts (seemingly mainly *Rhinolophus* bats)^13-16^. In our comparative genomic analysis, we found ORF3c to be the only novel ORF that is conserved across sarbecoviruses and subject to purifying selection^10^. Translation of ORF3c is supported by ribosome profiling data^1^, which is not the case for the majority of other predicted novel SARS-CoV-2 ORFs. ORF3c thus represents a previously uninvestigated area of sarbecovirus research.

SARS-CoV-2 infection is known to dysregulate host immune responses, specifically the type I interferon (IFN) response, resulting in the severe clinical symptoms characteristic of this pathogen^17^. Type I IFNs are essential innate cytokines that induce a host response that restricts and eliminates SARS-CoV-2 infection^18^. Host cells produce type I IFNs in response to activation of pattern recognition receptors (PRRs): host proteins that recognise molecules from either pathogens or damaged cells. Pathogen-associated molecular patterns (PAMPs) activate PRRs that in turn promote transcription of type I IFNs and other antiviral genes^19^. The two key cytosolic PRRs involved in the antiviral response are retinoic acid-inducible gene (RIG-I) and melanoma differentiation-associated protein 5 (MDA5). RIG-I senses short dsRNA and 5′-ppp/pp-RNA^20^ whereas MDA5 senses high molecular weight dsRNA and mRNA that lack 2′-O-methylation at the 5′ cap^21,22^. Both RIG-I and MDA5 form filaments with dsRNA to recruit mitochondrial antiviral-signalling protein (MAVS). This interaction promotes the formation of MAVS polymers at the mitochondrial outer membrane that activate transcription factors interferon regulatory factor 3 (IRF3) and nuclear factor kappa-light-chain-enhancer of activated B cells (NF-κB)^23^. The activated IRF3 dimer and NF-κB complex translocate into the nucleus and bind to responsive promoters to stimulate the transcription of type I IFNs^24^ and other pro-inflammatory factors. To suppress production of these molecules, viruses have evolved different strategies targeting the activation of RIG-I, MDA5 or MAVS. For example, multiple SARS-CoV-2 proteins directly and indirectly target MAVS to diminish IRF3 activation and so production of IFN-β and other pro-inflammatory proteins encoded by IRF3 responsive genes. Membrane (M) protein interacts with MAVS to suppress MAVS polymerisation, leading to diminished IFN-β production and enhanced SARS-CoV-2 replication^25^, whereas ORF9b interacts with mitochondrial import receptor subunit Tom70 at mitochondria to suppress MAVS and reduce activation of the IFN-β promoter^26^. ORF9b has also been found to bind to IKK-γ (NEMO), a protein that acts downstream of MAVS^27^. Finally, when ORF10 is overexpressed, it activates mitophagy receptors to induce MAVS elimination^28^.

Here we present an initial functional characterisation of ORF3c. We find that it localises to the mitochondrial outer membrane, a platform heavily involved in innate immune signalling. We provide evidence that ORF3c subverts the cascade of cellular antiviral responses via preventing activation of transcription from the IFN-β gene. This appears to be mediated by interactions of ORF3c with both PGAM5 (mitochondrial protein phosphoglycerate mutant family member 5, previously known as phosphoglycerate mutase family member 5) and MAVS, alongside a subsequent cleavage of MAVS by activated caspases.

## RESULTS

### ORF3c is conserved across sarbecoviruses

Since we identified ORF3c as a previously undetected and conserved ORF in 2020^10^, numerous additional sarbecovirus sequences have been published, including many divergent sequences from bat hosts. We inspected GenBank sequence records for all unique ORF3c sequences (Figure 1A, Figure S1). The ORF3c protein is 39–41 amino acids in length and has a predicted C-terminal transmembrane region and a shorter N-terminal hydrophobic region (Figure 1A).

**Figure 1.**
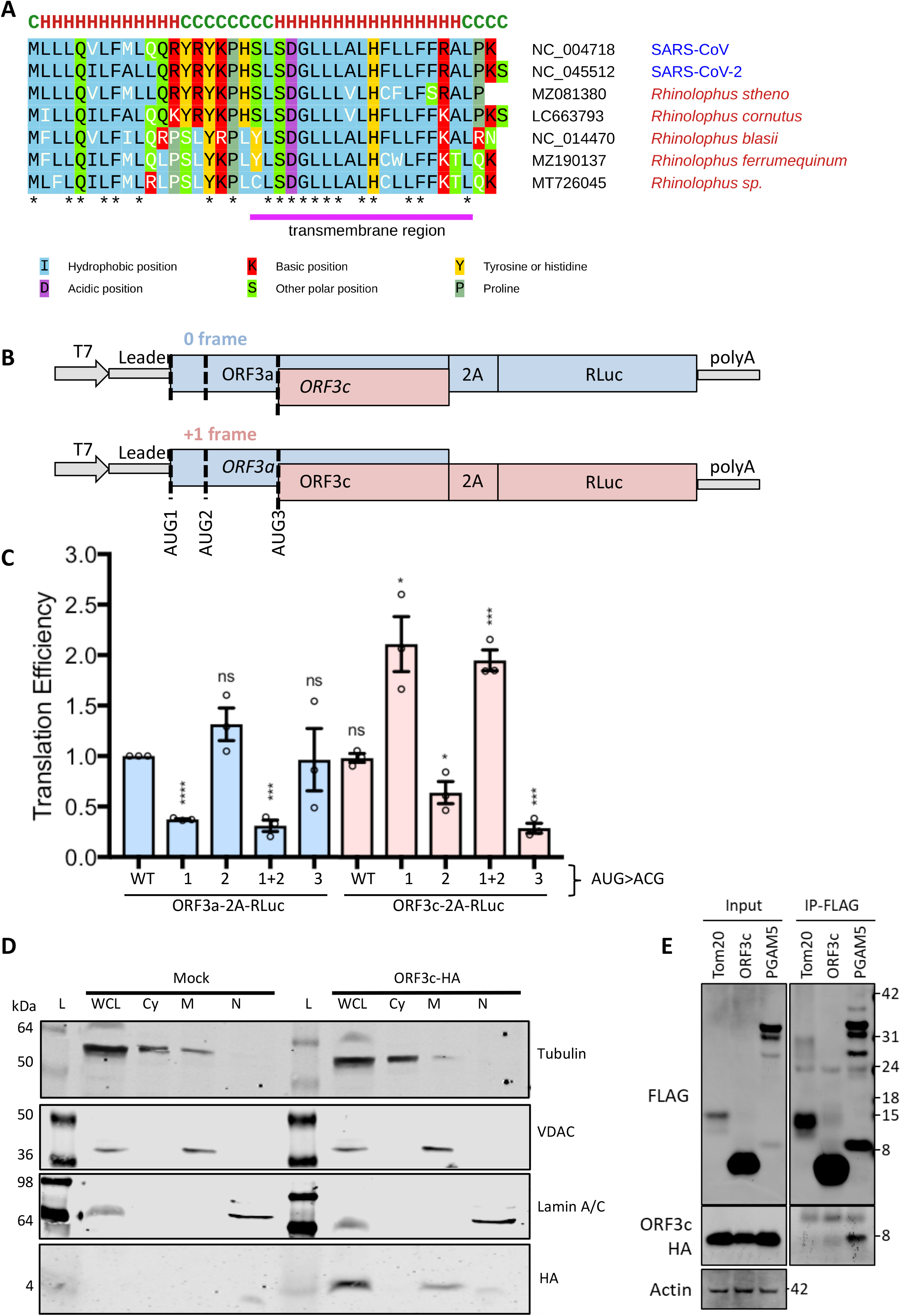
ORF3c is a conserved membrane-associated protein expressed by leaky scanning. (A) Representative ORF3c sequences from different sarbecoviruses. See Figure S1 for additional sequences. Sequences are labelled with accession numbers and either name of host (red) or name of virus (blue). Sequence differences from SARS-CoV-2 ORF3c are indicated with white letters. Amino acids are colour-coded according to their physicochemical properties. Asterisks indicate completely conserved columns in the alignment. A transmembrane region predicted by Phobius^133^ is indicated with a pink bar below the alignment. The helix(H)/coil(C) secondary structure of SARS-CoV-2 3c as predicted with RaptorX^134^ is shown above the alignment. (B) Schematic of *in vitro* transcribed reporter RNAs. RNAs contained the SARS-CoV-2 ORF3a sgmRNA from the leader to the end of the ORF3c coding region. This was followed by FMDV 2A and RLuc sequences, which were in-frame with one of the three AUG codons: two in the ORF3a frame (0 frame), and the last in the ORF3c frame (+1 frame). The AUGs were then mutated systematically to ACG to inhibit ribosomal initiation. (C) RLuc values were normalised to an internal FLuc control, for each well. Normalised values for each individual experiment were converted to a percentage of ORF3a WT. Individual data points (circles), and means (bars) ± SEM (error bars) based on 3 independent experiments are shown. Statistical analysis (unpaired two sample t-test): ns = not significant, *P ≤ 0.05, **P ≤ 0.01, ***P ≤ 0.001, ****P ≤ 0.0001. (E) Vero cells were transfected with ORF3c-HA or empty plasmid (Mock) and subjected to subcellular fractionation at 24 h post-transfection. Fractions were probed by immunoblot for various markers: alpha-tubulin (cytoplasmic), VDAC (mitochondrial membrane), lamin A/C (nuclear) and the HA tag. A HA-specific band was only observed in the membranous fraction. WCL: whole cell lysate; Cy: cytoplasmic fraction; M: membranous fraction; N: soluble nuclear fraction; L: protein markers (ladder). (E) ORF3c does not form multimeric complexes. HEK293T cells were transfected with plasmids expressing FLAG-tagged Tom20, ORF3c or PGAM5 and ORF3c-HA. The following day, cell lysates were collected and proteins were immunoprecipitated by the FLAG epitope. FLAG-tagged proteins and HA-tagged ORF3c of input (Left) or IP (Right) samples were detected by immunoblotting with rabbit anti-FLAG and mouse anti-HA antibodies.

### ORF3c is expressed via leaky scanning

ORF3c is a small protein (predicted MW = 4.9 kDa) and the ORF entirely overlaps with ORF3a in the +1 frame (with respect to ORF3a). We hypothesised that ORF3c is expressed via ribosomal leaky scanning on the ORF3a sgmRNA, in a similar manner to the ORF7b and ORF9b proteins of SARS-CoV-1^29,30^. This would require scanning preinitiation (43S) complexes to proceed past both the AUG start codon of ORF3a and a subsequent in-frame AUG, and then initiate at a third AUG: the start codon of ORF3c (mechanism reviewed in Firth, 2012^31^). Both AUGs in the ORF3a frame have intermediate or weak initiation contexts (thus facilitating leaky scanning) whereas the ORF3c AUG has a strong initiation context^32^.

To test the leaky scanning hypothesis and measure the ORF3a:ORF3c relative expression levels, an expression cassette was created composed of the 5′ end of the ORF3a sgmRNA transcript (including the 77 nucleotide leader) up to the final coding nucleotide of ORF3c (excluding the stop codon). This was followed by a foot and mouth disease virus (FMDV) 2A sequence (which mediates co-translational separation of the polypeptide chain) and then the Renilla luciferase (RLuc) sequence. The 2A-RLuc ORF was in-frame with the AUG of either ORF3a or ORF3c. A range of mutants were created based on both of these constructs, which ablated (i) the first AUG of ORF3a; (ii) the second AUG of ORF3a; (iii) both AUGs of ORF3a; or (iv) the AUG of ORF3c (Figure 1B). In all cases, AUGs were mutated to ACG (which, however, may still allow low level initiation^31^).

Plasmid DNA templates were transcribed *in vitro* by the T7 RNA polymerase and the transcripts were then purified and transfected into Vero cells in a 96-well plate format. The luciferase values were measured at 20 h post transfection. RLuc was normalised to an internal firefly luciferase (FLuc) control. Values for the ORF3c WT and ORF3a and ORF3c mutant RNAs were then normalised to those for the ORF3a WT RNAs (Figure 1C). Expression of ORF3a or ORF3c was sensitive to, respectively, mutation of the first or both ORF3a-frame AUGs, and mutation of the ORF3c-frame AUG. However, the second AUG was not noticeably utilised for ORF3a expression. As expected, mutation of the ORF3c-frame AUG did not affect ORF3a expression. With the WT sequence, the levels of ORF3c and ORF3a expression appeared to be similar to each other. Consistent with a leaky scanning model, when the first or both ORF3a-frame AUGs were mutated, ORF3c expression approximately doubled. Thus, under the conditions tested, approximately 50% of preinitiation complexes appear to scan past the first two AUGs within the ORF3a sgmRNA and initiate at the next downstream AUG to translate ORF3c. This translational efficiency would result in an approximately equal stoichiometric ratio of ORF3a:ORF3c proteins from the one sgmRNA.

### ORF3c associates with cellular membranes

Many mature SARS-CoV-2 proteins possess transmembrane domains or are membrane-associated via a range of topologies and anchors. These include the structural proteins spike (S), membrane (M) and envelope (E), non-structural proteins NSP4 and NSP6 and the accessory proteins ORF3a, ORF6, ORF7a, ORF7b and ORF9b. Other viral proteins possess N-terminal hydrophobic ER-targeting signals that are cleaved following membrane translocation and folding, resulting in a mature secretory-like protein (for example, ORF8). Membrane-tethered viral proteins localise to a range of subcellular locations: NSP3, NSP4 and NSP6 form a complex that rearranges the ER, where viral replication organelles are found^33,34^; ORF6 localises to the plasma membrane^35^, Golgi and ER^36^; ORF7a and ORF7b to the Golgi^37,38^; ORF9b to the mitochondrion and various intracellular vesicles (anchored notably via a hydrophobic cavity^39,40^, rather than a prototypic transmembrane helix); and ORF3a to lysosomes, Golgi^41^, ER and the plasma membrane^42,43^. The three transmembrane structural proteins S, M and E all localise with varying degrees to the ER, Golgi and ERGIC compartments during the viral assembly, budding and trafficking processes^44^.

ORF3c is predicted to possess a C-terminal □-helix (Figure 1A), which is likely to form a transmembrane domain^10^, suggesting that ORF3c is a single-pass integral transmembrane protein. To confirm this predicted membrane association, C-terminally HA-tagged ORF3c (ORF3c-HA) was overexpressed by transfection in Vero cells and the cells were subject to subcellular fractionation. Cytoplasmic, membranous and nuclear fractions were probed for ORF3c-HA via immunoblotting and ORF3c was found to localise primarily within the membranous fraction (Figure 1D).

### ORF3c does not multimerise

Many viruses encode similarly small transmembrane proteins that spontaneously multimerise to form ion channels within host membranes (so-called viroporins, reviewed in Nieva, 2012^45^). Previously, we and others suggested that ORF3c may possess viroporin activity due to its size and hydrophobic nature^10,11^. To assess the ability of ORF3c to multimerise, we co-transfected ORF3c-HA and ORF3c-FLAG into HEK293T cells and performed an anti-FLAG immunoprecipitation. FLAG-tagged PGAM5 was included as a positive control to ensure interaction partners were not lost (see later results and Figure 5). The eluate was then probed for both HA and FLAG tags by immunoblotting (Figure 1E). Whereas FLAG-tagged PGAM5 co-immunoprecipitated ORF3c-HA, ORF3c-FLAG did not. Thus, this experiment did not provide evidence for ORF3c homo-oligomerisation in cultured cells, suggesting that ORF3c is unlikely to form ion channels.

### ORF3c can localise to the ER membrane *in vitro*

According to the predicted C-terminal helical transmembrane domain^10^, ORF3c most likely is a tail-anchored protein. Whilst this group of diverse and functionally important integral membrane proteins are present in all intracellular membranes with a cytosolic surface^46,47^, it is generally accepted that most tail-anchored proteins bearing transmembrane domains of substantial hydrophobic character are post-translationally targeted to the endoplasmic reticulum (ER) – the central organelle involved in coronaviral RNA replication^48-50^ and to which multiple SARS-CoV-2 proteins localise. In contrast, tail-anchored proteins containing transmembrane domains of reduced hydrophobicity and increased charge within the extremely short exoplasmic C-terminal region (C_exo_), typically target to the mitochondrial outer membrane (MOM)^51,52^. Thus, we explored sequentially the ability of ORF3c to integrate into the membrane of the ER and mitochondria using *in vitro* systems specialised for each organelle.

Having exploited canine pancreatic microsomes to study the integration of other SARS-CoV-2 membrane proteins into the ER^53^, we studied ORF3c biogenesis using this system (Figure 2A). We found that *in vitro* synthesised and imported ORF3c (Figure 2B) was protected from added protease (Figure 2C, lanes 1-3) and was resistant to extraction with alkaline sodium carbonate buffer (Figure 2C, lanes 4-6), suggesting that ORF3c can integrate stably into the ER membrane *in vitro*.

**Figure 2.**
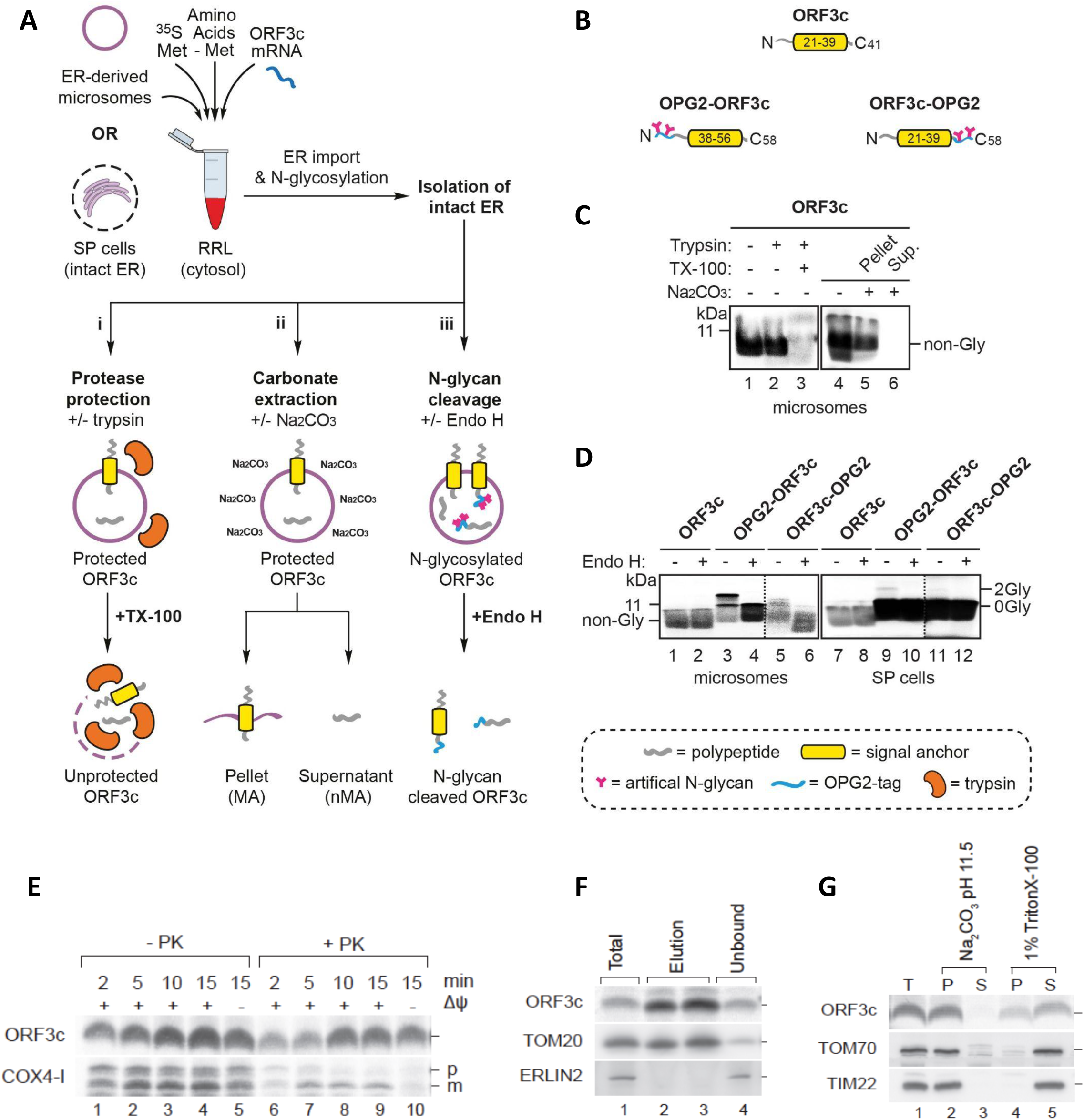
ORF3c can localise to the ER membrane *in vitro* but also inserts efficiently into mitochondrial membranes. (A) Outline of the *in vitro* assay where either canine pancreatic microsomes or semi-permeabilised (SP) HeLa cells are used as sources of ER membrane. Following translation, membrane-associated ORF3c proteins were recovered by centrifugation and subjected to: i) a protease protection assay using trypsin in the presence or absence of Triton X-100 (TX-100), ii) alkaline sodium-carbonate extraction, or iii) endoglycosidase H (Endo H) treatment. Resulting products were analysed by SDS-PAGE and phosphorimaging. (B) Schematics of parent ORF3c and its N-terminally (OPG2-ORF3c) and C-terminally (ORF3c-OPG2) OPG2-tagged variants. Comprising residues 1-18 of bovine rhodopsin (Uniprot: P02699), the OPG2 tag is indicated in blue and its N-glycan sites in pink. The putative ORF3c transmembrane domain is indicated in yellow. (C) Membrane-associated products of ORF3c synthesised in the presence of ER-derived microsomes were treated with trypsin (lanes 2-3) or sodium carbonate (lanes 5-6) as outlined in **Ai**-**Aii.** (D) Using ER-derived microsomes (lanes 1-6) or SP cells (lanes 7-12), parent and OPG2-tagged variants of ORF3c were synthesised as outlined in **Aiii**. N-glycosylated (2Gly) and non-glycosylated (0Gly) species were confirmed using Endo H. Note that, whilst on the same gel, the signals in lanes 5-6 and 11-12 of panel (D) have been overexposed compared to lanes 1-4 and 7-10 (demarcated by dotted lines) in order to enhance the visibility of the radiolabelled products. RRL, rabbit reticulocyte lysate. (E) *In vitro* import and proteinase K (PK) accessibility assay of isolated HEK293T mitochondria upon [^35^S]ORF3c-5M and COX4-1 import. p, precursor protein; m, mature protein. (F) Immuno-isolation of mitochondria with anti-Tom22 magnetic beads after [^35^S]ORF3c-5M import. (G) Chemical protein extraction following [^35^S]ORF3c-5M protein import into mitochondria isolated from HEK293T cells. Mitochondria were treated with sodium carbonate (Na_2_CO_3_, pH 11.5) or with TBS 1% Triton X-100. T, total; P, pellet; and S, soluble.

To investigate its membrane topology, we modified ORF3c to facilitate the detection of ER import – incorporating an OPG2 tag either at the extreme N- or C-terminus to generate OPG2-ORF3c and ORF3c-OPG2, respectively (Figure 2B). Since the OPG2 epitope supports efficient ER lumenal N-glycosylation^54^, we synthesised radiolabelled ORF3c and its two OPG2-tagged ORF3c variants in the presence of ER membranes and used endoglycosidase H (Endo H) treatment of the resulting membrane-associated products to identify N-glycosylated species (Figure 2D, even numbered lanes). On the basis of these studies, we found that both the N- and C-termini of OPG2-tagged ORF3c can be N-glycosylated and hence can be translocated into the lumen of ER microsomes (Figure 2D, lanes 1-6). Interestingly, when the same ORF3c proteins were analysed using semi-permeabilised HeLa cells (SP HeLa cells; Figure 2D, lanes 7-12), the extent of this N-glycosylation was greatly reduced but the total ORF3c signal increased, suggesting it may be targeted to another organelle.

Whereas canine pancreatic microsomes are highly enriched in ER-derived membranes, SP cells preserve the integrity of multiple subcellular organelles including both the ER and mitochondria^51,55^. Therefore, we considered the possibility that, in contrast to the robust N-glycosylation observed using purified ER membranes, the presence of mitochondria in SP cells may reduce the opportunity for ORF3c to mislocalise to the ER by providing access to the mitochondrial outer membrane (MOM)^51,56^.

### ORF3c inserts efficiently into mitochondrial membranes *in vitro*

Next, we investigated the ability of ORF3c to insert into the membranes of isolated mitochondria by using an *in vitro* assay comparable to that used to assess ER import. Tail anchored proteins are not found in the inner mitochondrial membrane, whereas a range of endogenous MOM proteins (such as Tom5, Tom6 and Tom7) possess this topology. We radiolabelled ORF3c in reticulocyte lysate and incubated the protein with mitochondria purified from HEK293T cells. ORF3c increasingly associated *in vitro* with mitochondria over time (Figure 2E). In contrast to the mitochondrial matrix protein COX4-1, association of ORF3c with mitochondria occurred independent of an inner mitochondrial membrane potential. Following ORF3c import, mitochondria were treated with proteinase K to degrade the non-imported protein. We observed a gradual increase in ORF3c signal intensity with time (Figure 2E, lanes 6-10) suggesting that the protein becomes proteinase K resistant probably due to membrane integration. To rule out that ORF3c targeting occurred due to a distinct contaminating-membrane in the mitochondrial preparation, anti-Tom22 antibodies immobilised on magnetic beads were used to immuno-isolate mitochondria after ORF3c import. The resulting immuno-isolated mitochondria showed clear enrichment for ORF3c and Tom20, while ERLIN2 (an ER marker) was not copurified (Figure 2F). Accordingly, we conclude that ORF3c is targeted selectively to mitochondria.

To address whether imported ORF3c was integrated stably into mitochondrial membranes, we imported ORF3c into purified mitochondria that subsequently were treated with sodium carbonate buffer (pH 11.4). Under these conditions, integral membrane proteins are retained in the membrane pellet, whilst soluble and peripheral membrane proteins are released into the supernatant. Upon carbonate treatment, ORF3c was present exclusively in the membrane pellet fraction, indicating that it was incorporated into the lipid phase (Figure 2G, lanes 2-3). Alternatively, mitochondrial membranes were solubilised in detergent (Triton X-100). Upon detergent treatment, ORF3c was largely released into the supernatant, although a fraction remained unsolubilised (Figure 2G, lanes 4-5). The residual amount of ORF3c recovered in the pellet fraction after Triton X-100 treatment indicated that ORF3c is prone to aggregation in the presence of the ionic detergent. In conclusion, our *in vitro* data strongly suggest that ORF3c is targeted to and inserted in mitochondrial membranes.

### ORF3c localises to the mitochondria in cells

To confirm the mitochondrial localisation of ORF3c in intact cells, Vero cells were transfected transiently with ORF3c-HA and immunofluorescence microscopy was performed to identify its subcellular localisation. Clear membrane staining was observed and cells were co-stained using antibodies against various sub-cellular markers. No co-localisation was observed between ORF3c-HA and tubulin, ER markers (calnexin, PDI), Golgi markers (RCAS1, TGN46), or endolysosomal markers (early: EEA1, late: CD63, lysosome: LAMP1). However, clear co-localisation was observed using antibodies against the MOM import receptors Tom20 and Tom70 (Figure 3A).

**Figure 3.**
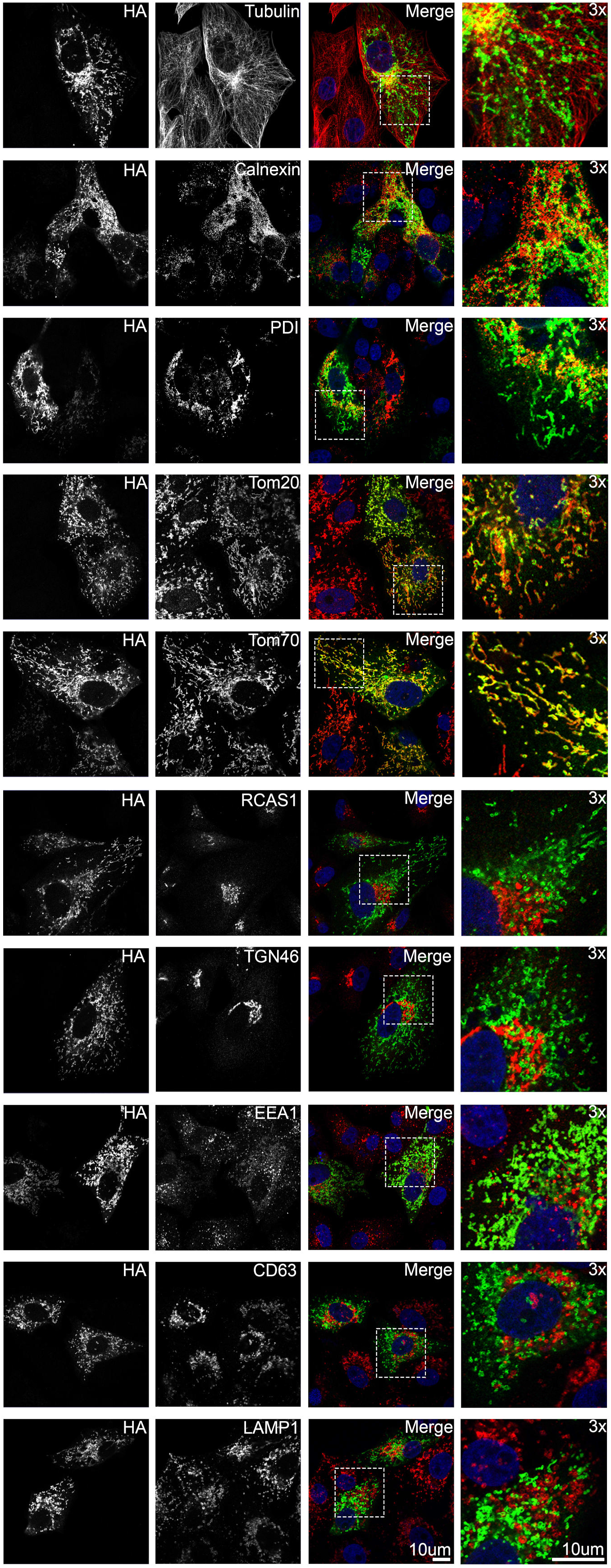
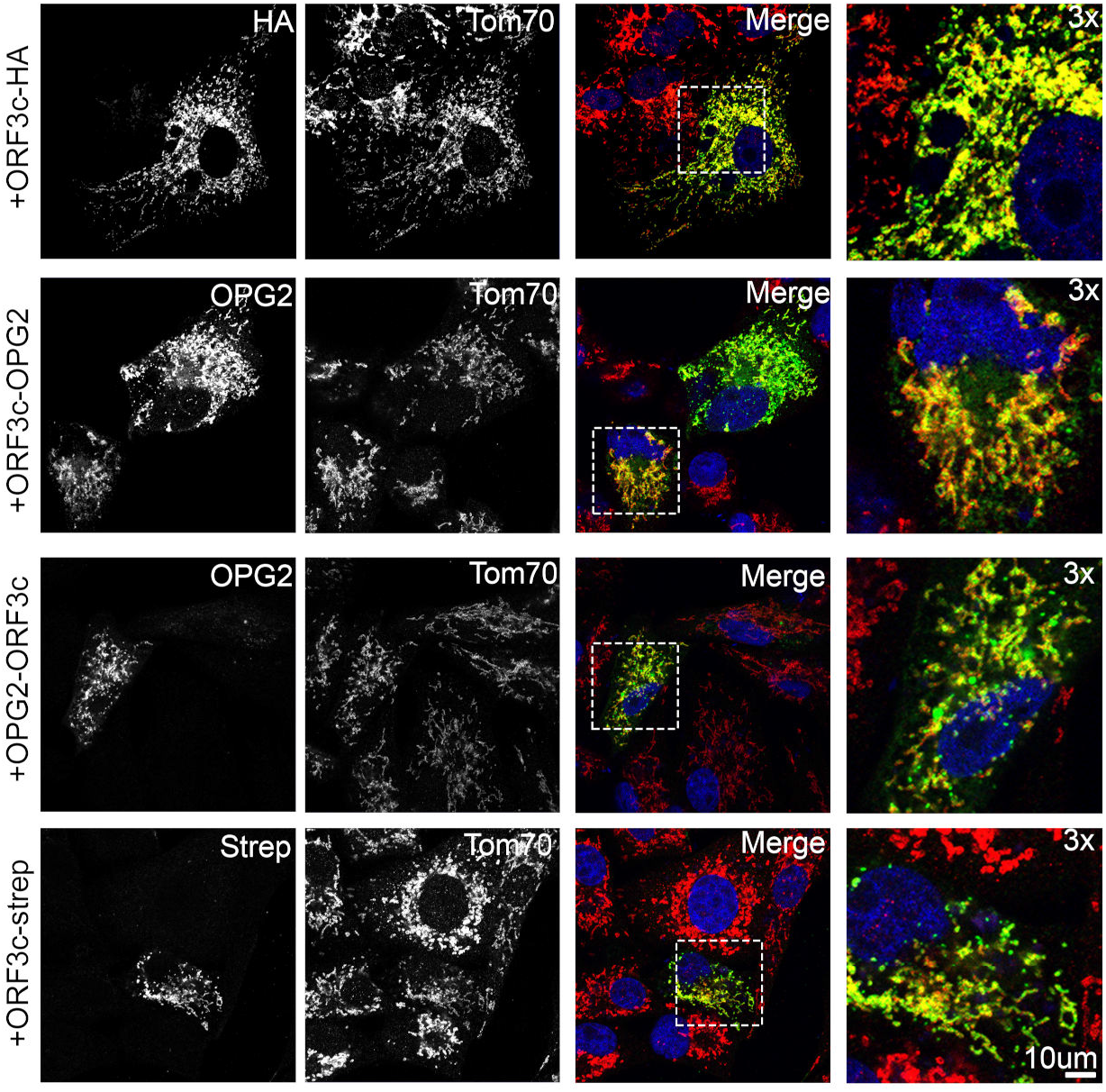
ORF3c localises to the mitochondria in cells. (A) Vero cells were transfected with a plasmid expressing ORF3c-HA and 24 h later the cells were fixed and co-stained using the indicated antibodies raised against a range of endomembrane marker proteins and against HA. (B) Vero cells were transfected with plasmids expressing alternatively tagged ORF3c proteins, including ORF3c-OPG2, OPG2-ORF3c and ORF3c-Strep. Fixed cells were co-labelled using antibodies against each epitope tag and against Tom70.

To ensure that the C-terminal HA tag was not mis-directing ORF3c-HA localisation, alternative epitope tags (OPG2 and Strep) were fused to either the N or C terminus of ORF3c. Ectopic expression of all ORF3c versions displayed clear co-localisation with endogenous Tom70 (Figure 3B). Furthermore, the mitochondrial localisation of ORF3c did not alter over 48 h post-transfection (Figure S2).

### PGAM5 interacts specifically with ORF3c

To identify host interaction partners, we immunoprecipitated ORF3c-HA from transfected Vero cells grown in labelled medium (stable isotope labelling of amino acids in culture, SILAC). ORF3c-HA-binding proteins were compared in triplicate to a HA-only control. Immunoprecipitated samples were subject to trypsin digest and then analysed by liquid chromatography with tandem mass spectrometry (LC-MS/MS).

Six proteins were identified as being enriched in ORF3c-HA samples relative to HA-only control samples with a p-value, q-value and FDR all < 0.05 (Figure S3), and minimum of 0.5 log2 (i.e. 1.41-fold) enrichment. These were: PGAM5, RPL8, EIF6, ARPC5, CAVIN1 and CAPZB. The full set of quantified filtered proteins, and list of significantly enriched proteins are included as supplementary files (Supplemental Table 1, Supplemental Table 2). Of these six proteins, PGAM5 was chosen for further investigation due to its recently reported role in antiviral signalling^57,58^. The ORF3c:PGAM5 interaction was confirmed by co-immunoprecipitation of PGAM-FLAG and ORF3c-HA (Figures 1E, 5B).

PGAM5 is a single pass transmembrane protein with a cytosolic serine/threonine phosphatase domain^59^, which localises to the MOM via its N-terminal transmembrane domain^60^. Here it oligomerises into dodecamers, the catalytically active state^61-63^. Besides roles in cell death related processes and mitochondrial dynamics, recently PGAM5 was found to play a role in upregulating IFN-β signalling during infection by viruses from multiple families^57,58^, acting via a direct interaction with MAVS – a tail-anchored protein that also localises to the MOM following oligomerisation^64,65^. PGAM5 multimerisation, induced as a result of viral or poly(I:C) stimulation, causes increased phosphorylation of TBK1 and IRF3 and a subsequent increased transcription of IRF3-responsive genes including IFN-β. Interestingly, this function of PGAM5 appears to be independent of its phosphatase activity, despite the dodecameric form of PGAM5 being catalytically active^57^. It has also been reported that SARS-CoV-2 infection results in increased ubiquitylation of PGAM5, accompanied by an overall decrease in PGAM5 protein levels, suggesting it is targeted to the proteasome during infection^66^. Therefore, we investigated the ability of ORF3c to dysregulate innate immune signalling in stimulated cells, hypothesising that the ORF3c:PGAM5 interaction would abrogate the PGAM5-driven antiviral effect.

### ORF3c inhibits IFN-β signalling

We co-transfected ORF3c with luciferase reporter plasmids, wherein luciferase expression is driven by a range of promoters responsive to innate immune signalling, into HEK293T cells that were stimulated with either TNF-□, PMA, Sendai virus (SeV) or IFN-□. These would stimulate the promoters responsive to, respectively, NF-κB, activator protein 1 (AP-1), IFN-β and ISG56.1, or the interferon-stimulated response element (ISRE). Similar approaches have been used extensively to identify SARS-CoV-2 immune antagonists^17,27,67-70^. The results indicated that the transcription from the IFN-β-responsive promoter, but not those responsive to NF-κB or ISRE, is inhibited by ORF3c (Figure 4A). This appears to be mediated by the AP-1 element within the IFN-β promoter (Figure 4A, 4B) in a dose-dependent manner (Figure S4A). Other coronaviral proteins, known to antagonise this pathway, were also included as controls (Figure S4B, S4C). Importantly, we observed the same trend regardless of the ORF3c tag position, or indeed using untagged ORF3c (Figure 4B). Intriguingly, we could not detect the N-terminal HA tag, indicating that the N terminus may possess an internal cleavage site (Figure 4B, see also Figure 6A). Finally, to decrease the probability of artefacts arising due to the system choice (a concern that has been raised in relation to other potential SARS-CoV-2 immune antagonists^71^), we performed qRT-PCR analysis of SeV-infected A549 cells expressing ORF3c-FLAG stably (Figure 4D). This indicated that the *IFN-β, ISG56* and *ISG54* mRNAs were specifically downregulated in ORF3c-expressing SeV-stimulated cells (Figure 4C), indicating this blockade is at the transcriptional level. The downregulation of *ISG56* mRNA level in qRT-PCR is inconsistent with the previous ISG56.1 reporter gene assay, which might be due to sensitivity differences between the two techniques, and between HEK293T and A549 cells. Given that A549 cells are derived from human alveolar cells and are broadly used in respiratory virus infection, the qRT-PCR data from these cells that analyses the transcription of endogenous genes is more compelling than the reporter gene assays. Collectively, these data provide evidence that ORF3c might contribute to the suppression of the IFN-β signalling pathway during SARS-CoV-2 infection in cultured cells.

**Figure 4.**
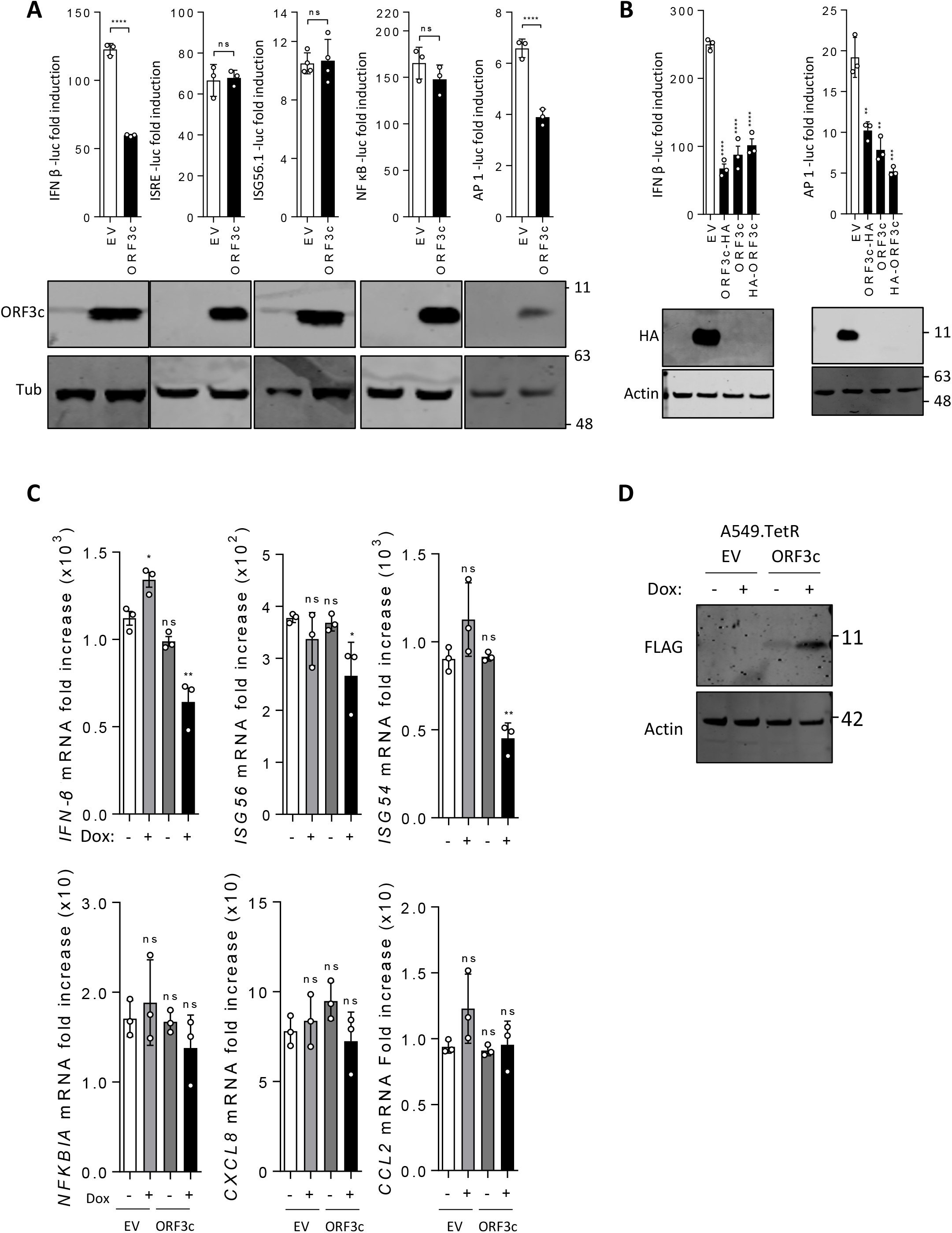
ORF3c inhibits IFN-β signalling. (A) HEK293T cells were co-transfected with plasmids encoding IFN-β, ISRE, NF-κB, ISG56.1 or AP-1-driven FLuc, together with RLuc and ORF3c-HA plasmids. On the following day, cells were stimulated with SeV (IFN-β and ISG56.1), 1,000 units/mL IFN-α (ISRE), 20 ng/mL TNF-α (NF-κB) or 20 ng/mL PMA (AP-1). SeV infection and PMA stimulation were maintained overnight whereas cytokine treatments were maintained for 6 h. The same stimulation conditions were used in all experiments unless otherwise stated. Cell lysates were collected after stimulation to measure luciferase levels and protein expression. FLuc activity was normalised to RLuc and the fold induction is shown relative to unstimulated controls. Below the figures are representative immunoblots of indicated protein expression. (B) The same reporter gene assays as described in (A) were performed with ORF3c tagged with HA at either the N or C terminus, or untagged ORF3c. (C) After overnight induction with tetracycline, ORF3c-inducible cells were infected with SeV for 6 h, and then collected to extract cellular mRNA. Total mRNA was reverse transcribed into cDNA and the indicated genes were quantified by qPCR. mRNA levels of target genes were normalised to the *GAPDH* mRNA level, and fold induction was calculated relative to the unstimulated control. Statistical analysis (unpaired two sample t-test): ns = not significant, *P ≤ 0.05, **P ≤ 0.01, ***P ≤ 0.001, ****P ≤ 0.0001. (D) Tetracycline-inducible A549 cells were induced or mock induced with 100 ng/mL doxycycline to express FLAG-tagged ORF3c.

**Figure 5.**
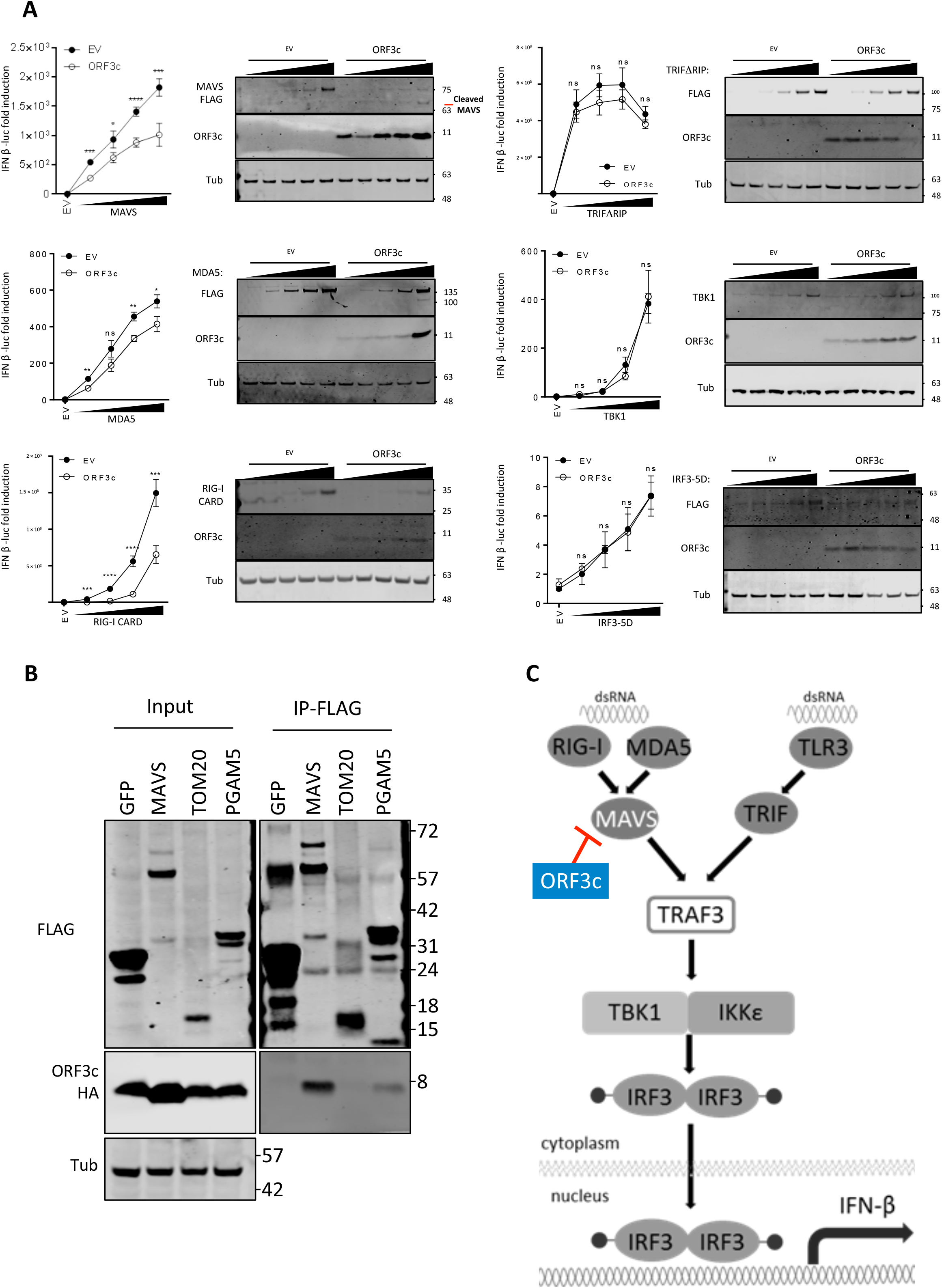
ORF3c interacts with MAVS to restrict IFN-β promoter activation. (A) HEK293T cells were co-transfected with plasmids expressing ORF3c or empty vector (EV), RLuc, and IFN-β-driven FLuc plasmids, together with increasing dose of plasmids expressing FLAG-tagged CARD domain of RIG-I (RIG-I-CARD), MDA5, MAVS, TRIFΔRIP, IRF3-5D or TBK1. Cell lysates were prepared and analysed as in Figure 4. Adjacent immunoblots show the expression of the indicated proteins. Statistical analysis (unpaired two sample t-test): ns = not significant, *P ≤ 0.05, **P ≤ 0.01, ***P ≤ 0.001, ****P ≤ 0.0001. (B) HEK293T cells were seeded in 10-cm dishes and transfected with 2 µg of plasmids encoding FLAG-tagged GFP, MAVS, Tom20 or PGAM5, and 2 µg of ORF3c-HA plasmid. After 48 h, cell lysates were collected, subjected to immunoprecipitation with anti-FLAG affinity beads, and analysed by SDS-PAGE and immunoblotting. (C) Schematic illustrating the simplified pathways leading from RNA sensing to activation of IRF3 and transcription of the IFN-β gene. The position at which ORF3c mediates inhibition is indicated.

**Figure 6.**
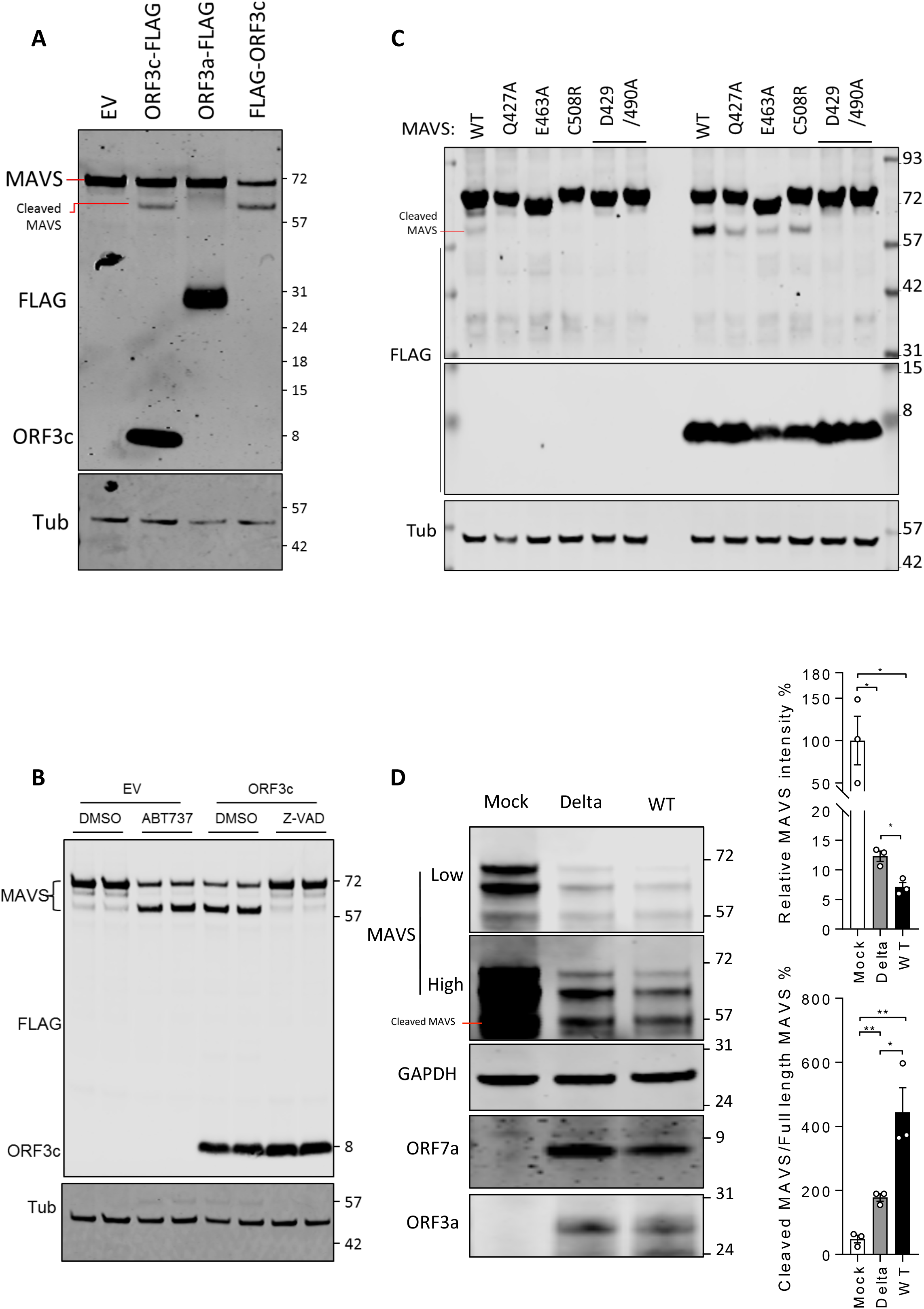
ORF3c induces MAVS cleavage. (A) HEK293T cells were co-transfected with empty vector (EV) or plasmids expressing FLAG-ORF3c, ORF3a-FLAG or ORF3c-FLAG and FLAG-MAVS. At 24 h post-transfection, cell lysates were collected and analysed by SDS-PAGE and immunoblotting with the antibodies indicated. Positions of full-length and cleaved MAVS are indicated by red lines. (B) Cells were transfected as described in (A) and treated with 20 µM ABT-737, 40 µM Z-VAD or DMSO overnight. Cell lysates were collected and analysed as described in (A). (C) HEK293T cells were co-transfected with plasmids expressing FLAG-tagged MAVS wild type (WT) or mutants Q427A, E463A, C508R or D429A/D490A, with (lanes 7-12) or without (lanes 1-6) ORF3c-FLAG. Cell lysates were collected at 24 h post transfection and processed as described in (A). (D) A549 +ACE2 +TMPRSS2 cells were infected with SARS-CoV-2 (Wuhan) or delta variant (EPI_ISL_1731019) at MOI 2. At 24 h post infection, infected and mocked-infected cells were collected and lysed. The cell lysates were analysed by immunoblotting using the indicated antibodies. MAVS immunoblots are shown at high and low intensity for clarity. The intensity of full-length MAVS (∼72 kDa) from 3 technical repeats was normalised to GAPDH, and then further normalised relative to mock infection and is shown in percentage (upper bar graph). The ratio of cleaved (∼57 kDa) / full length (∼72 kDa) MAVS was calculated and is shown in percentage (lower bar graph). Quantification of band intensity was performed with the LI-COR imaging system. Statistical analysis was based on three technical repeats of the same cell lysates (* P < 0.05).

### ORF3c interacts with MAVS to restrict IFN-β promoter activation

Next, we co-transfected FLAG-tagged proteins from within the IRF3-signalling pathway (RIG-I CARD, MDA5, MAVS, TRIFΔRIP, TBK1 or IRF3-5D^72^) alongside the IFN-β promoter-driven FLuc reporter and ORF3c. These proteins would activate the IFN-β promoter from various nodes and allow us to elucidate the stage(s) targeted by ORF3c. The results indicated that ORF3c acts upstream of the TRAF3/TBK1 nexus (Figure 5A). These data, alongside published results that PGAM5 interacts with MAVS^57^, led us to hypothesise that ORF3c interacts with MAVS. To investigate this, we performed a co-immunoprecipitation. Tom20 was included to assess whether we were isolating mitochondrial membrane proteins nonspecifically. The results indicate that both MAVS and PGAM5 are specific interaction partners of ORF3c (Figure 5B). Immunofluorescence experiments were also strongly supportive of co-localisation of ORF3c and MAVS in intact membranes (Figure S5). MAVS overexpression is sufficient to induce expression from the IFN-β and AP-1 responsive promoter^65,73^. Thus we hypothesise that sequestration of MAVS due to interaction with ORF3c may underlie the observed decrease in transcription from the IFN-β gene (Figure 5C).

### ORF3c induces MAVS cleavage

During these experiments, we noticed an additional, lower molecular weight specific band appearing in the MAVS immunoblots upon ORF3c co-transfection. This was reproducible but did not appear upon MAVS/ORF3a co-transfection, indicating this MAVS cleavage was an ORF3c-specific effect (Figure 6A). As it was inhibited by treatment with the pan-caspase inhibitor z-VAD, and mimicked the effects of apoptosis induction via the Bcl-2 inhibitor ABT737 (Figure 6B), this suggests that ORF3c-induced MAVS cleavage may be mediated by caspases and accompanied by apoptosis. To further investigate the ORF3c-mediated MAVS cleavage, we examined MAVS mutants that had reported resistance to viral protein or caspase-mediated cleavage^74-76^. We found that the MAVS point mutations D429A/D490A, but not Q427A, E463A or C508R, blocked the ORF3c-induced cleavage, indicating the cleavage is mediated by caspase-3 (Figure 6C).

Intriguingly, ORF3c is not expressed in the SARS-CoV-2 variant of concern delta (B.1.617.2) lineage due to a CAG to UAG mutation that introduces a premature termination codon at codon 5. The delta (B.1.617.2) variant therefore represents an opportunity to examine the effects of ORF3c depletion, albeit with the caveat that many other mutations co-exist in this genome compared to earlier lineages. We utilised this naturally occurring difference to examine MAVS levels in infected cells (Figure 6D). In these experiments the relative intensities of the three MAVS-related bands differed from the previous experiment (Figure 6A), possibly as a result of the different cell type (A549 versus HEK293T) or detection of endogenous rather than FLAG-tagged MAVS. Nonetheless, we found that MAVS is severely downregulated in infected cells compared to mock, and both the downregulation and the ratio of cleaved MAVS to full-length MAVS were somewhat reduced in delta-infected cells compared to cells infected with an early SARS-CoV-2 lineage possessing an intact ORF3c.

### ORF3c presence does not alter mitochondrial morphology in infected cells

Given the lack of ORF3c expression in the delta (B.1.617.2) variant, we next examined the mitochondrial morphology in A549 +ACE2 +TMPRSS2 cells infected with either delta (B.1.617.2-Por1, B.1.617.2-Por2) or non-delta (VIC1 and B.1.1.7) SARS-CoV-2 variants. Immunofluorescence staining for mitochondrial markers (Tom20 and Tom70) (Figure 7A, Figure S6A) indicated that, although mitochondrial staining was reduced in infected cells compared to uninfected cells, no clear differences were apparent between the variants. Additionally, by transmission electron microscopy, no phenotypic differences in the mitochondria of cells infected with different variants were observed (Figure 7B). This is perhaps unsurprising, due to the multiple redundant mechanisms of IFN dysregulation conferred by different SARS-CoV-2 proteins.

**Figure 7.**
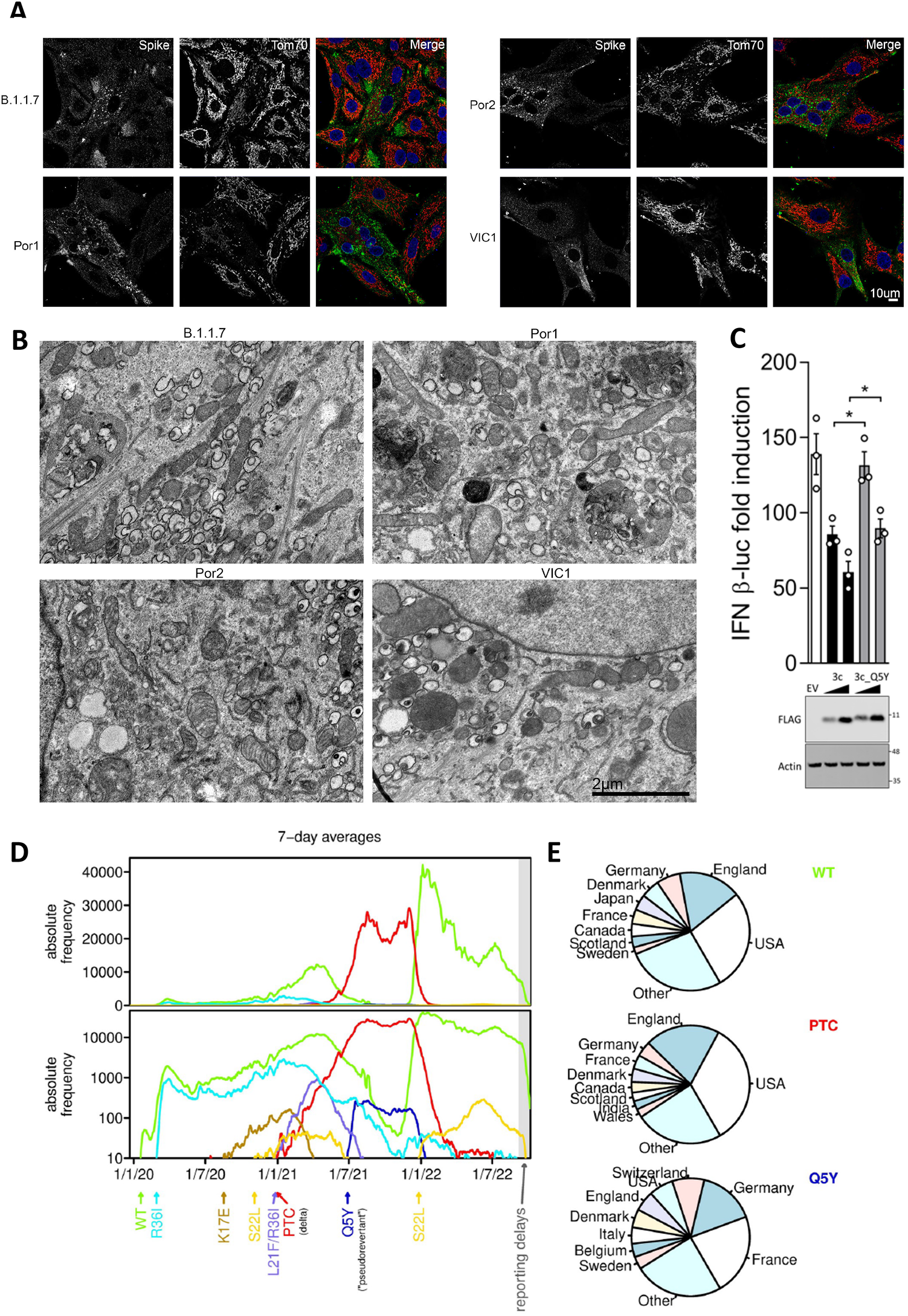
ORF3c absence does not alter mitochondrial morphology in infected cells. (A) A549 +ACE2 +TMPRSS2 cells were infected with either early lineage (VIC1, B.1.1.7) or delta lineage (B.1.617.2-POR1, B.1.617.2-POR2) isolates and fixed at 24 h post-infection. Cells were stained using antibodies against Spike and Tom70 and imaged by confocal microscopy. (B) As for (A) but cells were processed for electron microscopy imaging. Infected cells were identified by the presence of double membrane vesicles. (C) HEK293T cells were transfected with plasmids encoding RLuc, IFN-β driven FLuc, and two doses of either WT ORF3c-HA or a Q5Y mutant. The cells were stimulated, and cell lysates were collected and analysed as described in Figure 4A. (D) SARS-CoV-2 sequences present in the GISAID database that had specific-day collection dates specified were analysed. Data for all ORF3c variants present in 0.1% or more of the 12,906,225 ORF3c sequences analysed are shown. The two graphs show the same data, but on linear (upper panel) and log (lower panel) scales. The PTC mutant (red curve) corresponds to the SARS-CoV-2 delta variant. (E) Pie charts showing the geographic distribution of sequences obtained with the WT ORF3c 5th codon (CAG), PTC mutant 5th codon (UAG) and Q5Y “pseudorevertant” 5th codon (UAU).

### SARS-CoV-2 ORF3c variants in the human host

As noted above, ORF3c coding capacity is lost in the SARS-CoV-2 delta (B.1.617.2) variant, due to the appearance of a premature termination codon. To further study the appearance and dynamics of different ORF3c variants we queried 12,906,225 SARS-CoV-2 sequences from the GISAID database^77^ with coverage of the ORF3c region. Seven ORF3c amino acid variants were present at an abundance of ≥0.1% of the total, namely the original variant (WT), besides R36I, L21F/R36I, S22L, Q5Y, K17E and the aforementioned delta premature termination codon variant (PTC) (Figure 7D, Figure S6B). The CAG to UAG substitution at codon 5 of ORF3c that gave rise to the PTC variant pseudoreverted to UAU in the Q5Y variant. This restored expression of full-length ORF3c but with a Q to Y amino acid change at position 5 (the overlapping ORF3a amino acids are SD in WT, LD in PTC, and LY in Q5Y). This Q5Y pseudorevertant increased rapidly in mainland Europe sequencing reports around July 2021 before levelling off and eventually dying away as the delta virus variant was replaced with the omicron virus variant (B.1.1.529) in late 2021 (Figure 7E). Reporter gene assays indicated that the ORF3c Q5Y pseudorevertant was less efficient than the WT at antagonising IFN-β production but still had a marked effect (Figure 7C).

## DISCUSSION

Coronaviruses encode a variety of accessory proteins in their genomes, which are non-essential for RNA replication but confer advantageous properties to the virus allowing efficient viral propagation in the host. Many of these accessory proteins are known antagonists of the innate immune response and are useful targets for antiviral treatment strategies. However their variability across the *Coronaviridae* often means that studies cannot be extrapolated to other members; for example, SARS-CoV-1 ORF3b, which overlaps the 3′ region of ORF3a, is truncated in SARS-CoV-2^5^; and ORF10 in SARS-CoV-2 is entirely lacking in SARS-CoV-1. Previously, using comparative genomics, we identified ORF3c, an accessory protein conserved across the *Sarbecovirus* subgenus^10^. Here we have presented a functional analysis of ORF3c, revealing it to be a tail-anchored transmembrane protein that appears to be inserted into the mitochondrial outer membrane, where it interacts with MAVS and PGAM5, and reduces IFN-β signalling.

To the best of our knowledge, this is the first report of PGAM5 being involved in coronaviral-host protein:protein interactions, although it had been identified as a potential target for viral-induced proteasomal degradation^66^. PGAM5 localises to the MOM^60^, although there are also reports of PGAM5 within the inner mitochondrial membrane with the C-terminal catalytic domain facing the intermembrane space^78,79^. It has been suggested that its location may depend on cellular stress levels: PGAM5 is known to activate the MAP kinase pathway by dephosphorylating ASK1 (associated with cellular stress)^80^ and is involved in numerous cell death-related processes. Primarily, it is thought to regulate mitochondrial dynamics (fusion *versus* fission). Its ability to promote or suppress various cell death pathways is a contentious issue (recently reviewed in Cheng, 2021^81^): whilst reported initially as a pro-necrotic factor^59,82,83^, this has been disputed^84^. Equally controversial are its role(s) in apoptosis: it has been reported to suppress apoptosis in certain models^85^, yet be essential for apoptosis induction in others^86-88^. What does appear consistent – and dependent on the phosphatase function of PGAM5 – is a role in the induction of mitophagy, an organelle-specific form of autophagy that protects the cell against necroptosis by selectively degrading damaged mitochondria^87,89-92^.

It is unlikely that ORF3c affects the phosphatase function of PGAM5, given this is thought to be independent of the role of PGAM5 in immune signalling. PGAM5 multimerisation has been shown to be required for both IFN-β upregulation and induction of many cell death-related events. One possibility is that the ORF3c-induced decrease in IFN-β may be at least partially caused by a reduction in PGAM5 multimerisation due to its interaction with ORF3c, and a subsequent redirection of PGAM5 to the proteasome. Equally, it is possible that PGAM5 multimerisation continues in the presence of ORF3c, but the PGAM5:MAVS interaction is ablated. A third hypothesis involves the formation of a potential trimeric complex (ORF3c:MAVS:PGAM5), resulting in the functional abrogation of both host proteins. As the self-multimerisation of MAVS and PGAM5 are independent events^57^, this would be an efficient mechanism of sequestering both potential innate response activators with a single viral protein.

In addition to the observed ORF3c:MAVS and ORF3c:PGAM5 interactions that may inhibit PGAM5:MAVS stimulation of IFN-β production, we also observed cleavage of MAVS when ORF3c was overexpressed and this cleavage appeared to be driven by caspase-3 suggesting a link to apoptosis. Whether ORF3c-driven apoptosis is an artefact of overexpression outside of the context of virus infection (where other viral proteins might inhibit apoptosis) is currently unknown. While we found a tantalising suggestion of differences in MAVS cleavage between delta and non-delta infections, the overall strong downregulation of full-length MAVS in either infection compared to mock and the presence of cleaved MAVS even in mock-infected cells in this system, besides potential effects of other differences between delta and non-delta viruses, make it difficult to draw robust conclusions at this stage. Intriguingly, ORF3a of both SARS-CoV-1 and SARS-CoV-2, which localises to the plasma membrane, has been implicated in apoptosis induction; however in the case of SARS-CoV-1 this was mapped to the cytosolic C-terminal domain and therefore could not have been an incorrectly attributed function of ORF3c^93,94^.

PGAM5 has been reported to be cleaved (within the N-terminal transmembrane domain) in response to mitochondrial dysfunction and mitophagy, specifically during outer membrane rupture, resulting in its release into the cytosol^78,79,95^. Although we did not observe cleavage of PGAM5 in the presence of ORF3c (indicating mitophagy is not occurring), it would be of interest to confirm this with other methods. Equally, it would be of interest to analyse the phosphorylation patterns of PGAM5 when bound to ORF3c given that the phosphorylation status of this protein has numerous effects upon the downstream signalling pathways that it activates. During preparation of this manuscript, data became publicly available indicating ORF3c overexpression does not affect mitophagy, despite its mitochondrial localisation, but rather blocked autophagy by causing autophagosome accumulation^96^. This finding supports our preliminary evidence that ORF3c may direct cells towards apoptosis in preference to other cell death pathways, possibly by sequestration of PGAM5 and prevention of mitophagy activation.

Among SARS-CoV-2 proteins, ORF3c is not alone in having an inhibitory effect on IFN-β expression; however the mode of action differs significantly between the proteins involved. Some proteins directly reduce IFN-β mRNA or protein levels: ORF6, ORF8 and N all have similar effects to ORF3c and reduce IFN-β mRNA (and hence protein) levels although, unlike ORF3c, ORF6 and ORF8 simultaneously reduce expression from ISRE-containing promoters^67,68,97^. ORF6 has also been shown independently to reduce IRF3 and STAT1 nuclear translocation^17,36,69,98^; in comparison, N is thought to inhibit the TRIM25:RIG-I interaction^99,100^. NSP13^6,69^ and NSP6^69^ bind TBK1 directly, preventing IRF3 phosphorylation. Other sarbecoviral proteins inhibit type I IFN activation as an indirect result of their enzymatic function: SARS-CoV-1 NSP16 reduces MDA5 and IFIT activation by capping the viral RNA^101^; NSP14 reduces IFN levels by shutting down host translation^102^. Still others (NSP1 and NSP6) suppress the signalling induced by type I IFN, whilst leaving protein levels unaffected^69,103^. The convergent effects of these viral antagonistic proteins, which collectively target multiple layers of the immune signalling cascade, no doubt combine to reduce the host antiviral response and increase virus fitness in the natural host.

Despite the redundancy in IFN antagonists, those that operate directly from a mitochondrial location are uncommon amongst characterised coronaviral proteins. ORF9b and ORF10 are the exceptions. Similar to our observations for ORF3c, these proteins localise to the mitochondria upon overexpression and are able to dampen the immune response in the absence of other coronaviral proteins^28,40,104,105^, indicating they may each act in an unassisted fashion and not from within a virally encoded protein complex. ORF9b does, however, interact with host Tom70^6,26,104,105^ which in turn is known to interact with MAVS^106,107^. It has been suggested that the ORF9b:Tom70 interaction may lead to either apoptosis or mitophagy, as the levels of functional Tom70 will affect both of these processes; but these hypotheses have not been validated experimentally^105^. This provides an interesting parallel to ORF3c, which appears to operate at the same cellular location as Tom70 (specifically, the MOM^108^), yet we did not observe ORF3c and Tom70 to co-immunoprecipitate (data not shown). It remains possible that an indirect, transient interaction may occur between ORF9b, ORF3c, Tom70 and MAVS. Additionally, unlike ORF3c, ORF9b has been observed to inhibit the IKK-γ (NEMO) cascade^27^, suggesting that ORF9b has additional functions downstream of MAVS, specifically inhibiting the NF-κB pathway. Thus it appears that sarbecoviruses have evolved complementary approaches, mediated by ORF9b and ORF3c respectively, to subvert IFN-β signal transduction and reduce mitochondrial innate immune pathway activation from within the mitochondrion itself. The mechanism employed by ORF10 is different yet again; however this protein is not conserved across the subgenus. ORF10 localises to the mitochondrion where it interacts with the mitophagy receptor NIX, to activate mitophagy and thereby eliminate aggregated MAVS^28^. This may in part explain why downregulation and degradation of MAVS is still visible during infection with a delta variant lacking ORF3c (Figure 6D). Downregulation of MAVS has also been reported from proteome-wide studies of SARS-CoV-2 infected cells (although the level of reduction appears to depend on the model system)^109,110^.

Prior to the identification of ORF3c, a screen of viral and host protein:protein interactions did not identify either MAVS or PGAM5 as probable interaction partners for any SARS-CoV-2 protein^6^. This was reflected in a thorough literature review^111^. Equally, there are remarkably few confirmed direct interactions of MAVS with SARS-CoV-2 proteins. Although there are some reports of the M protein interacting with MAVS^25,70^, this is not reconciled with the lack of a mitochondrial localisation for the M protein which is found consistently at the ER and Golgi^70,105^. As such, the relevance of this potential interaction during an actual viral infection is open to question. In short, ORF3c is the only conserved sarbecoviral protein that has been shown to bind directly to MAVS within the MOM, where the majority of activated MAVS would be located during a viral infection.

There are several obstacles currently impeding further progress. For example, analysis of ORF3c-HA transfected cell lysates following digestion with trypsin or chymotrypsin and LC-MS/MS analysis failed to identify ORF3c-derived peptides even when inclusion lists of predicted ORF3c peptides were employed, likely due to high hydrophobicity of the peptides. This explains why multiple published analyses using trypsin digestion have failed to identify the ORF3c protein during infection (data not shown)^9,112^. Equally, our own work has been limited by the poor immunogenicity of ORF3c: a peptide-raised rabbit antibody was not reactive against transfected cell lysates, nor was a sheep polyclonal antibody raised against the entire ORF3c protein (data not shown). However, we are confident that these hurdles will be overcome with time.

The discovery of ORF3c necessitates a reassessment of previous sarbecoviral ORF3a-targeted studies, which may have also included ORF3c during protein overexpression. DNA-based constructs (although useful and often necessary) create an artificial system, because these vectors generally exclude viral untranslated regions and are designed to optimise expression from the desired AUG codon. Thus the degree of ORF3c expression, alongside the desired ORF3a, remains an unknown factor for most previous studies. It is also probable that ORF3c-mediated effects were overlooked in historical SARS-CoV-1 studies due to the extensive use of Vero cells, which allow efficient replication of many coronaviruses but are deficient in type I IFN production^113^. For example, inadvertent deletion of ORF3c via ORF3a mutation results in only minor attenuation in these cells, as measured by viral infectious titre^114,115^ (although, notably, deletion of the entire ORF3a region reduced cytopathic effect and cell death^116^).

Sarbecoviruses appear to be primarily bat viruses (Figure S1) and ORF3c appears to be conserved throughout this clade (with the exception of two different sarbecovirus sequences from *Rhinolophus hipposideros* where ORF3c is truncated; Figure S1B). In the human host, ORF3c is clearly not essential (as demonstrated by the success of the delta variant where ORF3c is disrupted). However, whether or not ORF3c provides a selective advantage in the human host is unclear. It is possible that the observed loss, restoration and variations of ORF3c in the human host may be random events whose effects on virus fitness are outweighed by increases in virus fitness conferred by other mutations (e.g. in the spike protein) with ORF3c variations being “carried along”. The importance of ORF3c in the human host presumably will become clearer as the virus adapts to long-term persistence in the human population. Similarly, other hosts such as the palm civet *Paguma larvata* and the Malayan pangolin *Manis javanica* may be intermediate hosts to which these viruses have not fully adapted. The observed Q5Y pseudorevertant of the delta PTC truncation is curious and may reflect a selective advantage of restoring ORF3c protein expression, but it might also have been a random event. Notably Q5 is perfectly conserved across the sarbecovirus ORF3c sequences (Figure S1A) suggesting that a Q at this position is functionally important, at least in bats. Sarbecoviruses have many different ways to antagonise host innate immunity and it may be that ORF3c is redundant in the human host (or its relative importance may also depend on cell type, host genetic background or disease state). Although not essential, ORF3c may still lead to an increase in virus fitness in the human host. Future work will be needed to compare WT and ORF3c knockout viruses in both human and bat cell lines, besides animal models.

## ACKNOWLEDGMENTS

The authors would like to thank Dr James Hastie and the staff at the MRC Protein Phosphorylation and Ubiquitylation Unit, University of Dundee, for providing anti-3c custom sheep antibodies. We would also like to thank the UK Health Security Agency for providing virus isolates. We gratefully acknowledge all data contributors, i.e. the Authors and their Originating laboratories responsible for obtaining the specimens, and their Submitting laboratories for generating the genetic sequence and metadata and sharing via the GISAID Initiative, on which some of this research is based.

## AUTHOR CONTRIBUTIONS

H.S., Y.L., A.E.F., G.L.S., J.R.E., S.H. and P.R. designed the study and experiments. H.S., Y.L., S.O’K., G.W.C., R.M., J.R.E., E.E., S.K.N., N.L., H.A.M., A.V., L.D.C.Z, D.A.M., and A.D.D. performed experiments. H.S., Y.L., S.H., J.R.E., E.E., V.L., I.J. and P.R. provided expertise on data analysis, interpretation and methodology. A.E.F. performed bioinformatic analyses. H.S., Y.L., and S.O’K. wrote the manuscript. All authors edited and discussed the manuscript. A.E.F., E.E., G.L.S. and J.L.H. secured funding.

## METHODS

### Cell culture and transfections

Vero (ATCC, CRL-1586), HEK293T (ATCC, CRL-3216), A549 cells (ATCC, CCL-185) and HeLa (ATCC, CCL-2) cells were maintained in high-glucose Dulbecco’s-modified Eagle’s medium (DMEM, Invitrogen) supplemented with 10% fetal bovine serum (FBS), 100 IU/ml penicillin, 100 μg/ml streptomycin, 1 mM nonessential amino acids, 25 mM HEPES, and 1% L-glutamine (complete DMEM) in a humidified incubator at 37°C with 5% CO_2_. A549 cells stably expressing ACE2 and TMPRSS2 (A549 +ACE2 +TMPRSS2) were kindly provided by Prof. Massimo Palmarini (University of Glasgow)^117^ and were maintained in the above-described medium, with 2 mg/mL G418 and 200 μg/mL hygromycin B. All cells were routinely tested and confirmed to be free of mycoplasma (MycoAlert™ PLUS Assay, Lonza).

For DNA transfections, cells were seeded in 6-well plates at 30–40% confluency the day prior to transfection. Four μg of pure plasmid DNA was transfected with 30 μl of Lipofectamine 2000 (Invitrogen) in a total volume of 1 ml serum-free OptiMem (Invitrogen) according to manufacturer’s instructions. At 3 h post-transfection, 1.5 ml of DMEM with 10% FBS was added to each well. Cells were harvested 24 h post-transfection.

HEK293T-Flp-InTM T-RexTM (HEK293T) cells were maintained and cultured in DMEM media (GiboTM, Thermo Fisher Scientific, Waltham, MA, USA) supplemented with 10% sterile filtered FBS (Sigma, Saint Louis, MO, USA) at 37°C with 5% CO_2_. For all the mitochondrial import experiments, HEK293T cells were freshly plated three days before to reach 80% confluency.

### Viruses

SARS-CoV-2 strains BetaCoV/Australia/VIC01/2020 (B lineage)^118^, BetaCoV/England/ MIG457/2020 (B.1.1.7), and delta variants ASL516 (B.1.617.2-POR1) and ASL517 (B.1.617.2-POR2) were obtained from Porton Down, UK Health Security Agency, and propagated in Vero +TMPRSS2 +hACE2 cells under BSL-3 conditions. SARS-CoV-2 delta variant (GISAID accession number: EPI_ISL_1731019) was kindly provided by Prof. Wendy Barclay. SARS-CoV-2 (Wuhan) (GenbankAccession MW041156.1) was isolated at the University of Bristol.

### Generation of tagged ORF3c expression constructs

The coding sequence of ORF3c fused with Strep or HA coding sequences (either N- or C-terminal) was amplified via PCR and each amplicon was inserted into the pCAG-PM vector^119,120^ using *Afl*II and *Pac*I restriction sites. A GGSGGS linker was used in these cases. The resulting plasmids were confirmed by Sanger sequencing (University of Cambridge, Department of Biochemistry DNA Sequencing Facility).

OPG2-tagged ORF3c plasmids were generated by site-directed mutagenesis (Stratagene QuikChange, Agilent Technologies) and confirmed by DNA sequencing (GATC, Eurofins Genomics). To improve the signal intensity of *in vitro* synthesised radiolabelled proteins, linear DNA templates containing an additional five methionine residues (5M) at the C terminus were generated by PCR, and linear DNA templates were transcribed into mRNA using T7 polymerase (Promega).

FLAG-tagged and untagged ORF3c expression vectors were constructed by PCR amplification from pCAG.ORF3a and inserted into pCAG-FLAG or pcDNA6B-FLAG. FLAG-tagged ORF3c mutant Q5Y was prepared by site-directed mutagenesis using a commercial kit (New England Biolabs, E0554S).

### Additional plasmids

Dual-luciferase reporter assay plasmids containing ISRE, NF-κB, AP-1 or IFN-β responsive promoters driving FLuc expression, and the plasmid expressing RLuc were described^121^. The ISG56.1 reporter plasmid was kindly provided by Dr Ganes Sen, Cleveland Clinic, Cleveland, OH. Vectors expressing MAVS, TBK1, TRIFΔRIP and IRF3-5D were described^121^. Vectors expressing RIG-I-CARD and MDA5 were provided by Dr Carlos Maluquer de Motes (School of Biosciences and Medicine, University of Surrey, UK) and Prof. Richard Randall (School of Biology, University of St Andrews, UK), respectively. Plasmids expressing MAVS mutants were generated by site-directed mutagenesis. SARS-CoV-2 vectors expressing ORF3a, NSP6, NSP9, NSP12 and empty vector pCAG.FLAG were kindly provided by Dr Peihui Wang, Shandong University, China^122^. PGAM5-FLAG-pcDNA3.1 and TOMM70A-FLAG-pcDNA3.1 constructs were purchased directly from GenScript (GenScript, Netherlands).

### Generation of an inducible ORF3c-expressing cell line

Inducible A549 cells expressing ORF3c were prepared by lentivirus transduction. The lentivirus preparation and infection have been described^123^. In brief, A549 cells were transduced with a lentivirus vector expressing a tetracycline repressor (TetR) linked to a nuclear localisation signal (NLS) and enhanced green fluorescent protein (EGFPnlsTetR). The transduced cells were selected with medium supplemented with 0.5 mg/mL G418, and then the EGFPnlsTetR positive cells were sorted by fluorescence activated cell sorting (FACS). The sorted cells were transduced with a second lentivirus vector expressing FLAG-tagged ORF3c. Transduced cells were then selected by medium supplemented with 500 ng/mL puromycin. The lentivirus vectors pLKO.neo.EGFPnls-TetR and pLKO.DCMV-TetO.mcs were kind gifts from Prof. Roger Everett, MRC University of Glasgow Centre for Virus Research, Glasgow, UK. The pLKO.DCMV.TetO.ORF3c plasmid was constructed in the G.L.S. laboratory.

### Reporter gene assays

HEK293T cells were seeded in 96-well plates with 3 × 10^5^ cells per well. After overnight incubation, cells were transfected with 100 ng/well of FLuc plasmids (ISRE, NF-κB, AP-1, IFN-β or ISG56.1), 10 ng/well of RLuc plasmid, and the indicated viral protein expression vectors. At 24 h post-transfection, the cells were stimulated as follows: 6 h stimulation of 1000 unit/ml IFN-□ (Peprotech, 300-02AA) for ISRE-luc, 6 h stimulation of 20 ng/ml TNF-□ (Peprotech, 300-01A) for NF-κB-luc, 24 h infection of SeV (Cantell Strain, Licence No. ITIMP17.0612A) for ISG56.1-luc and IFN β-luc, and 24 h stimulation of 10 ng/ml PMA (Sigma-Aldrich, P8139) for AP-1-luc. Stimulated, or infected, cells were lysed with 50 μL/well of passive lysis buffer (Promega) and kept at −20•. To measure luciferase activity, 50 μL FLuc reagent, or 50 μL RLuc reagent was added to 8 μL cell lysate. The luminescent value of the lysate was measured with a microplate reader (BMG Labtech). Relative luminescent values were calculated by normalisation of the FLuc value to the RLuc value, for each well. Sendai virus (SeV) used in this study was kindly provided by Professor Steve Goodbourn, St George’s Hospital Medical School, University of London.

### RT-qPCR

Inducible A549 cells expressing ORF3c-FLAG or empty vectors were seeded in triplicate at 4 × 10^5^ cells per well on 24-well plates. The next day, the cells were mock-induced or induced with 100 ng/mL doxycycline overnight. Then, the cells were infected with SeV for 6 h to stimulate IFN-β transcription. The cells were harvested and total RNA was extracted using RNeasy Mini Kit (Qiagen, 74106). Thereafter, 500 ng RNA was used in cDNA synthesis using SuperScript III reverse transcriptase (Invitrogen, 18080093). The expression of genes of interest was analysed by qPCR. Each cDNA sample was duplicated in the reaction, using SYBR Green Master Mix (Thermo Fisher Scientific, 4309155). A ViiA 7 real-time PCR system (Thermo Fisher Scientific) was used to determine each reaction cycle threshold. Amplification of the investigated genes was normalised to GAPDH amplification. Fold induction of each gene was calculated relative to unstimulated control of each condition.

### Leaky scanning assays

A plasmid was engineered in which a T7 promoter sequence preceded the first 77 nucleotides (the leader sequence) of the ORF3a subgenomic RNA. This was followed by the ORF3a coding region, up to but excluding the stop codon of ORF3c. This was followed by a foot and mouth disease virus 2A peptide sequence and then a RLuc ORF. The RLuc was in-frame with the AUG (start) codon of either ORF3a or ORF3c; the plasmids were termed pL_3a_SGRluc and pL_3c_SGRluc respectively. These plasmids were linearised with FastDigest *Aan*I (Invitrogen) and RNA was *in vitro* transcribed using the T7 mMessage kit (Invitrogen) and terminated with DNase treatment. RNA was purified with a silica-gel based membrane method (RNA Clean and Concentrator, Zymo Research) and the integrity was confirmed by gel electrophoresis. FLuc RNA was used as a transfection control. This RNA was generated by T7 *in vitro* transcription, using an in-house FLuc plasmid linearised with *Bam*HI as described^124^.

RNA transfections were conducted in triplicate in 96-well plates. Vero cells were seeded the day prior at 40% confluency (1 × 10^5^ cells per well). One hundred and fifty ng of L3a_SGRluc RNA (or that of a derived mutant) plus 30 ng of FLuc RNA was transfected into each well using 1 μl of Lipofectamine 2000, according to the manufacturer’s instructions, in a total volume of 40 μl of sera-free OptiMem. At 3 h post-transfection, 60 μl of DMEM with 10% FBS was added to each well. Cells were harvested at 20 h post-transfection with passive lysis buffer (Promega) and a dual luciferase detection assay was conducted upon the lysates (Promega).

### SDS-PAGE and immunoblots

Lysates from transfected cells were analysed by sodium dodecyl sulphate polyacrylamide gel electrophoresis (SDS-PAGE) using Tris-Glycine, at the percentage polyacrylamide suitable for resolution of the protein of interest. Precast Novex 10–20% tricine protein gels (Thermofisher Scientific) were used to resolve proteins under 10 kDa. Proteins were transferred to 0.2 μm nitrocellulose membranes and blocked with a commercial blocking buffer (LI-COR) before being probed with the relevant primary and secondary antibodies. Membranes were imaged using an Odyssey CLx imaging system (LI-COR). Primary antibodies used in this study were anti-FLAG (1:1000, rabbit polyclonal, SIGMA, F7425), anti-HA (1:500, mouse monoclonal, BioLegend, 901501), anti-actin (1:1000, rabbit polyclonal, SIGMA, A2066), anti-ORF7a (1:500, sheep polyclonal, University of Dundee, MRC PPU) and anti-ORF3a (1:500, sheep polyclonal, University of Dundee, MRC PPU). Secondary antibodies used in this study were anti-mouse IgG (1:10,000, donkey, LI-COR, 926-68072), anti-mouse IgG (1:10,000, goat, LI-COR, 926-32210), anti-Rabbit IgG (1:10,000, donkey, LI-COR, 926-68073), anti-mouse IgG (1:10,000, goat, LI-COR, 926-32211) and anti-goat IgG (1:10,000, donkey, LI-COR, 926-68074).

### Subcellular fractionation

Vero cells were transfected with a range of plasmids as described above. At 24 h post-transfection, monolayers were washed in phosphate-buffered saline (PBS) and subject to subcellular fractionation using a commercially available kit (Thermofisher Scientific) according to the manufacturer’s instructions. An input sample (10%) was retained for analysis prior to the first lysis step. Target-specific antibodies were anti-HA (Abcam, ab20084, 1:1000), anti-alpha-tubulin (Abcam, ab15568, 1:1000), anti-VDAC1 (Abcam, ab14734, 1:1000), and anti-lamin A/C (Abcam, ab133256, 1:1000).

### Immunoprecipitation

To detect protein-protein interactions, HEK293T cells (3 × 10^6^ cells per plate) were transfected with 2 μg of FLAG-tagged GFP, MAVS, TOMM70, TOMM20 or PGAM5 plasmids and 2 μg of ORF3c-HA expressing plasmid. At 48 h post transfection, transfected cells were washed with ice-cold PBS and lysed with cell lysis buffer (200 mM NaCl, 50 mM Tris, 1% NP40, pH 7.0) supplemented with cOmplete Mini EDTA-free protease inhibitor cocktail (Roche, 11836170001). The lysates were cleared by centrifugation at 13,000 rpm, and the FLAG-tagged proteins were immunoprecipitated with anti-FLAG agarose (A2220, MERCK). Input samples were collected as described above.

### Confocal microscopy

Vero cells were transfected transiently with ORF3c plasmids (described above). At 24 h post-transfection, cells were trypsinised and re-seeded upon sterile 13 mm glass coverslips. At 48 h post-transfection, the cell culture medium was removed and coverslips were washed once with PBS prior to fixation with 4% PFA/PBS.

For MitoTracker staining, the stock solution of MitoTracker® Red CM-H2XRos (Invitrogen, M7513) was prepared with DMSO to a concentration of 1 mM. Before staining live cells on coverslips, MitoTracker was diluted to 100 nM with pre-warmed DMEM without serum. The cells were washed twice with pre-warmed DMEM to remove serum and then incubated with MitoTracker containing medium at 37 °C for 1 h before fixation.

For SARS-CoV-2 infections, A549 +ACE2 +TMPRSS2 cells were grown on glass-bottomed 24-well plates (MatTek, P24G-0-13-F). Cells were infected at an MOI of 1 and placed on a rocker at room temperature for 2 h. The inocula were removed and replaced with full medium. At 24 h post infection plates were submerged in 4% PFA/PBS for 20 min to fix cells.

Free aldehydes were quenched with 15 mM glycine/PBS before cells were permeabilised with 0.1% saponin/PBS. Blocking and subsequent steps were performed using 1% BSA/0.01% saponin/PBS. Coverslips were inverted into droplets of primary antibody for 1 h, before being washed three times with PBS. Coverslips were then placed onto a second droplet containing fluorophore-conjugated secondary antibody for 45 min. Coverslips were washed three times before being mounted on to glass slides using ProLong Gold Antifade mountant containing DAPI (Invitrogen, P36931). Cells were imaged using a LSM700 confocal microscope (63x/1.4 NA oil immersion objective; ZEISS).

Primary antibodies used were anti-Tom20 (1:200, mouse monoclonal, Abcam, ab283317), anti-Tom70 (1:500, rabbit polyclonal, Proteintech, 14528-1-AP), anti-Calnexin, (1:250, rabbit monoclonal, Cell Signalling Technology, C5C9), anti-RCAS1 (1:250, rabbit monoclonal, Cell Signalling Technology, D2B6N), anti-Syntaxin 6 (1:250, rabbit monoclonal, Cell Signalling Technology, C34B2), anti-alpha-tubulin (1:500, mouse monoclonal, ProteinTec, 66031-1), anti-EEA1 (1:500, mouse monoclonal, BD Biosciences, 14/EEA1), anti-CD63 (1:400, mouse monoclonal, BioLegend, H5C6), anti-LAMP1 (1:400, mouse monoclonal, BioLegend, H4A3), anti-TGN46 (1:500, rabbit polyclonal, abcam, ab50595), anti-HA (1:500, rabbit polyclonal, Cell Signalling Technology, C29F4), anti-HA (1:500, rat monoclonal, Roche, 3F10), anti-OPG2 (1:100, mouse monoclonal^125^), StrepMAB-Classic (1:500, mouse monoclonal, IBA lifesciences, 2-1507-001), anti-SARS-CoV-2 Spike (1:100, mouse monoclonal, GeneTex, 1A9), anti-MAVS (1:100, rabbit polyclonal, Cell Signalling Technology, 3993), anti-ORF3a (1:100, sheep polyclonal, University of Dundee, MRC PPU), anti-HA (1:500, mouse monoclonal, BioLegend, 901501), anti-FLAG (1:500, rabbit polyclonal, Sigma, F2555).

Secondary antibodies used in this study were Alexa Fluor 488 goat anti-rat IgG (1:400, Invitrogen, A-11006), Alexa Fluor 488 goat anti-mouse IgG (1:400, Invitrogen, A-11001), Alexa Fluor 488 donkey anti-sheep IgG (A-11015), Alexa Fluor Plus 555 goat anti-rabbit IgG (1:400, Invitrogen, A32732).

### Electron microscopy

Cells were seeded upon plastic Thermanox coverslips in 24-well plates. Following infection, plates containing infected cells (see virus infections methods) were submerged in 2% PFA / 2.5% glutaraldehyde / 0.1 M cacodylate buffer, pH 7.4. Cells were washed with 0.1 M cacodylate buffer before being stained using 1% osmium tetroxide:1.5% potassium ferrocyanide for 1 h, and staining was further enhanced with UA-Zero (Agar Scientific) for 30 min. Cells were washed, dehydrated and infiltrated with Epoxy propane (CY212 Epoxy resin:propylene oxide) before being embedded in Epoxy resin. Epoxy was polymerised at 65°C overnight before Thermanox coverslips were removed using a heat-block. Seventy nm sections were cut using a Diatome diamond knife mounted to an ultramicrotome. Ultrathin sections were stained with lead citrate. An FEI Tecnai transmission electron microscope at an operating voltage of 80 kV was used to visualise samples.

### Semi-permeabilised (SP) cell preparation

HeLa cells (human epithelial cervix carcinoma cells, mycoplasma-free), as described^53,54,126^, were provided by Martin Lowe (University of Manchester) and were cultured in DMEM supplemented with 10% (v/v) FBS (Gibco, 10500-064) and maintained in a 5% CO_2_ humidified incubator at 37°C. Cells were seeded at 1 × 10^6^ per 10 cm^2^ dish and, once ∼80% confluent, cells were semi-permeabilised using digitonin (Calbiochem) and endogenous mRNA was removed by treatment with 0.2 U Nuclease S7 Micrococcal nuclease, from *Staphylococcus aureus* (Sigma-Aldrich, 10107921001) and 1 mM CaCl_2_ as described^53,54^. After quenching by the addition of EGTA to 4 mM final concentration, SP cells were resuspended in an appropriate volume of KHM buffer (110 mM KOAc, 2 mM Mg(OAc)_2_, 20 mM HEPES-KOH pH 7.2) to give a suspension of 3 × 10^6^ SP cells/mL as determined by trypan blue staining (Sigma-Aldrich, T8154). Freshly prepared SP cells were then included in translation master mixes such that each translation reaction contained 2 × 10^5^ cells/mL.

### *In vitro* ER import assays

Translation and membrane insertion assays, supplemented with nuclease-treated canine pancreatic microsomes (from stock with OD_280_ = 44/mL) or SP HeLa cells, were performed in nuclease-treated rabbit reticulocyte lysate (Promega) as described^53,54^. Briefly, in the presence of EasyTag EXPRESS ^35^S Protein Labelling Mix containing [^35^S] methionine (Perkin Elmer) (0.533 MBq; 30.15 TBq/mmol), 25 μM amino acids minus methionine (Promega), 6.5% (v/v) ER-derived microsomes or SP cells and 10% (v/v) of *in vitro* transcribed ORF3c mRNA (∼1 μg/μL) encoding the relevant ORF3c precursor protein. All translation reactions (30 μL) were performed at 30□C for 1 h and finished by incubating with 0.1 mM puromycin for 10 min at 30□C to ensure translation termination and the ribosomal release of newly synthesised proteins prior to analysis.

### Recovery and analysis of radiolabelled products synthesised in ER import assays

Following puromycin treatment, microsomal or SP cell membrane associated fractions were recovered by centrifugation through an 80 μL high-salt cushion [0.75 M sucrose, 0.5 M KOAc, 5 mM Mg(OAc)_2_ and 50 mM HEPES-KOH, pH 7.9] at 100,000 g for 10 min at 4°C. Then, the pellet was suspended in SDS sample buffer and, where indicated, samples were treated with 1000 U of endoglycosidase Hf (New England Biolabs, P0703S). To confirm association of ORF3c with the ER membrane, microsomal membrane-associated fractions were resuspended in KHM buffer (20 μL) and subjected to a protease protection assay using trypsin (1 μg/mL) with and without 0.1% Triton X-100 or sodium carbonate extraction (0.1 M Na_2_CO_3_, pH 11.3) as described^53^, prior to suspension in SDS sample buffer. All samples were solubilised for 12 h at 37°C prior to resolution by SDS–PAGE (16% PAGE, 120 V, 120 min). Gels were fixed for 5 min (20% MeOH, 10% AcOH), dried for 2 h at 65°C, and radiolabelled products were visualised using a Typhoon FLA-700 (GE Healthcare) following exposure to a phosphorimaging plate for 24–72 h.

### Mitochondria isolation

HEK293T cells were harvested in PBS, washed, and resuspended in 1X THE buffer (300 mM trehalose, 10 mM KCl, 10 mM HEPES, 1 mM EDTA, and 2 mM PMSF). Homogenization was performed in a PTFE pestle/glass Potter-Elvehjem at 700 rpm. The resulting cell lysate was centrifuged at 400 x *g* for 10 min at 4°C, followed by a second centrifugation step at 800 x *g* for 10 min at 4°C to remove unbroken cells. To sediment mitochondria, the supernatant was centrifuged at 10,000 x *g* for 10 min at 4°C. The mitochondrial sediment was washed with THE buffer and resuspended in the same buffer. The concentration of mitochondria was determined using a standard Bradford assay.

### *In vitro* import of [^35^S]ORF3c-5M into HEK293T isolated mitochondria

The ORF3c gene was amplified by PCR, adding five methionine codons to the C terminus (ORF3c-5M). The corresponding mRNA was prepared using the SP6 mMESSAGE mMACHINE kit (Invitrogen) according to the manufacturer’s specifications. Radiolabelled [^35^S]ORF3c-5M was synthesised *in vitro* in Flexi Rabbit Reticulocyte Lysate system (Promega) in the presence of 0.5 mM of KCl, 0.5 mM MgAc and 200 ng of ORF3c mRNA. For protein import, 100 µg of mitochondria resuspended in import buffer (250 mM sucrose, 80 mM potassium acetate, 5 mM magnesium acetate, 5 mM methionine, 10 mM sodium succinate, and 20 mM HEPES pH 7.4) with 1 mg/ml final concentration were used per reaction. [^35^S]ORF3c-5M protein was added at 5% (v/v) to the mitochondrial suspension and incubated at 37°C for 30 min under mild agitation (450 rpm). After import, mitochondria were sedimented at 10,000 x *g* for 10 min at 4°C and washed with HS buffer (500 mM sucrose and 20 mM HEPES/KOH pH 7.4).

### Proteinase K accessibility assay

After [^35^S]ORF3c-5M import, mitochondria were resuspended in TBS (intact mitochondria) or TBS + 1% Triton X-100 (lysed mitochondria), followed by digestion with proteinase K (PK) (10 µg/mL final concentration) for 10 min on ice. Next, PK was inhibited with 2 mM PMSF. Intact mitochondria were then washed. Proteins were precipitated with trichloroacetic acid (TCA). Samples were analysed by Tris/tricine SDS-PAGE. Proteins were transferred onto PVDF membranes. The [^35^S]ORF3c-5M protein signal was detected by digital autoradiography and mitochondrial proteins were immunodetected.

### Chemical extraction of mitochondrial membranes

After [^35^S]ORF3c-5M import, mitochondria were incubated in sodium carbonate buffer (pH 11.5) or in 1% Triton X-100 at 1 mg/mL final concentration for 20 min on ice. Insoluble material was sedimented at 100,000 x *g* for 1 h at 4°C. Pellet and soluble fractions were collected, and the protein was precipitated with TCA. Proteins were resolved on urea SDS-PAGE, transferred to PVDF membranes, and analysed for the [^35^S]ORF3c-5M protein by digital autoradiography and mitochondrial proteins by immunodetection.

### Immuno-isolation of mitochondria after ORF3c import

After import of [^35^S]ORF3c-5M, mitochondria were re-isolated with an anti-TOM22 mitochondria isolation kit (Miltenyi Biotec). In brief, resuspended mitochondria were incubated with anti-TOM22 Microbeads for 1 h with end-to-end mixing at 4°C. Beads were then collected through a MACS column placed in a magnetic MACS separator. The column was then washed to remove non-specific binding. After removing the column from the magnetic MACS separator, the beads were recovered and treated with TCA. Samples were separated on Tris/tricine SDS-PAGE, transferred onto PVDF membranes, and analysed by autoradiography to detect [^35^S]ORF3c-5M protein and immunodetection for the required proteins.

### SILAC immunoprecipitation experiment

Vero cells were grown in high glucose DMEM lacking arginine and lysine (Life Technologies), with 10% dialysed (7 kDa MWCO) FBS and supplemented with light (R0K0), medium (R6K4) or heavy (R10K8) stable isotope labelled arginine and lysine. Cells were maintained in labelled medium for 6 passages before transfection and immunoprecipitation. Labels were switched for the different replicates to control for any impact of the different culture media on cell growth/gene expression as follows: replicate 1 (L: HA-3c, M: Control), replicate 2 (M: HA-3c, H: Control) and replicate 3 (H: HA-3c, L: Control).

Vero cells were transfected with ORF3c-HA as described above. At 24 h post-transfection, monolayers were washed in ice-cold PBS and co-immunoprecipitation of the HA-tagged protein and interacting proteins was performed using a commercially available kit (HA Tag Magnetic IP/Co-IP kit, Pierce). An input sample (10%) was retained for analysis by immunoblot prior to binding the lysates to the magnetic anti-HA beads. After removing the remains of the wash buffer (third and final wash), samples were heated to 95• for 5 min in buffer consisting of 200 mM HEPES pH 8, 1% SDS and 1% NP40. Beads were collected by centrifugation and the supernatant was retained. Equal volumes of the control and HA-3c samples for each replicate were then combined and the samples were reduced, alkylated and trypsin-digested using the SP3 method^127^.

LC-MS/MS analysis was conducted on a Dionex 3000 coupled in line to a Q-Exactive-HF mass spectrometer. Digests were loaded onto a trap column (Acclaim PepMap 100, 2□cm□×□75□µm inner diameter, C18, 3 µm, 100 °A) at 5□µL per min in 0.1% (v/v) TFA and 2% (v/v) acetonitrile. After 3□min, the trap column was set in line with an analytical column (Easy-Spray PepMap® RSLC 15□×□50□cm inner diameter, C18, 2 µm, 100 °A) (Dionex). Peptides were loaded in 0.1% (v/v) formic acid and eluted with a linear gradient of 3.8–50% buffer B (HPLC grade acetonitrile 80% (v/v) with 0.1% (v/v) formic acid) over 95□min at 300□nL per min, followed by a washing step (5□min at 99% solvent B) and an equilibration step (25□min at 3.8% solvent). The Q-Exactive-HF was operated in data-dependent mode with survey scans acquired at a resolution of 60,000 at 200□*m*/*z* over a scan range of 350–2000□*m*/*z*. The top 16 most abundant ions with charge states +2 to +5 from the survey scan were selected for MS2 analysis at 60,000□*m*/*z* resolution with an isolation window of 0.7□*m*/*z*, with a (N)CE of 30%. The maximum injection times were 100 and 90□ms for MS1 and MS2, respectively, and AGC targets were 3e6 and 1e5, respectively. Dynamic exclusion (20□s) was enabled.

Data analysis was conducted in MaxQuant 1.6.7.0^128^. Options were set at default unless specified. Multiplicity was set at 3, with Arg6 and Lys4 set as ‘medium’ labels and Arg10 and Lys8 set as ‘heavy’ labels. Digestion was Trypsin/P, permitting up to two missed cleavages. Oxidation (M) and N-terminal protein acetylation were selected as variable modifications, and carbamidomethylation as a fixed modification. Under instrument settings, intensity was set to ‘total sum’. Fasta files containing the Uniprot African Green Monkey proteome (*Chlorocebus sabeus*, 19,223 entries, downloaded 16 May 2020) and a custom .fasta file containing HA-3c protein sequences were used for the search databases. The proteomics data generated in this study have been deposited in the ProteomeXchange Consortium (http://proteomecentral.proteomexchange.org) via the PRIDE repository (PXD037765).

Downstream data analysis was conducted in Matlab R2019a from the MaxQuant proteinGroups output file. Reverse database hits and common contaminants (MaxQuant contaminant list) were removed. Within each replicate, data were first normalised for equal protein loading based on the median SILAC intensity of shared proteins. Rows with >3 missing values were removed. Missing data were imputed (knn), and data were converted to HA-3C/HA-only ratios for each replicate, and then re-normalised across the three replicates by dividing by column median. A one-sample t-test was used to determine statistical significance. Multiple hypothesis testing was controlled using the approach of Storey, 2002^129^, with all meeting a FDR < 0.05. The Matlab scripts used for all data processing and figure generation with this data are available from the Emmott lab Github page at: https://github.com/emmottlab/sars2_3c/.

### Virus infections

To prepare lysates for immunoblots, A549 +ACE2 +TMPRSS2 cells were infected with SARS-CoV-2 (Wuhan) [GenBankAcc: MW041156.1] or delta variant [GISAID accession number: EPI_ISL_1731019, kindly provided by Prof. Wendy Barclay, Imperial College London] at a MOI of 2, or mock infected. At 24 h post-infection, cells were lysed directly with 2x SDS sample buffer and heated to 95• for 10 min. To prepare samples for electron microscopy, A549 +ACE2 +TMPRSS2 cells were grown on 13-mm Thermanox (plastic) coverslips (Nunc) in the wells of a 24-well plate. Cells were infected at an MOI of 1 and placed on a rocker at room temperature for 2 h. The inocula were removed and replaced with full medium and cells were returned to incubators for 24 h, at which point the plate was prepared for EM (see Electron microscopy methods).

### Identification of sarbecovirus ORF3c variants

To identify sequences with coverage of the ORF3a region, NCBI online tblastn^130^ was used on 11 Oct 2022, using the SARS-CoV-2 ORF3a protein (GenBank: YP_009724391.1) as query and the NCBI nr/nt database as subject, with organism taxonomy limited to *Coronaviridae* (taxid:11118) excluding SARS-CoV-2 (taxid:2697049), word size = 2, max target sequences = 1000, no low-complexity masking, and other parameters as defaults. The highest e-value of the 417 returned sequences was 8 × 10^−4^. The coordinates of tblastn matches on the subject sequences were extended to maximal stop-codon-to-stop-codon open reading frames, which were extracted, translated, and aligned as amino acid sequences with MUSCLE v3.8.31^131^. The amino acid alignment was used to guide alignment of the nucleotide sequences (EMBOSS:tranalign^132^) and the nucleotide sequences were 5′-truncated to the conserved ORF3a AUG initiation site. Sequences with defective or truncated ORF3a due to incomplete sequencing, frame-disrupting insertion/deletion errors, or other inconsistencies were discarded. An ORF homologous to SARS-CoV-2 ORF3c was not identified in the following sequences: MZ293757 (Hipposideros bat coronavirus, unclassified *Coronaviridae* member), and HQ166910 and NC_025217/KF636752 (subgenus *Hibecovirus*) and these sequences were therefore discarded. All remaining sequences had ≥62.6% amino acid identity to SARS-CoV-2 in ORF3a and may be regarded as members of subgenus *Sarbecovirus*, whereas the four discarded highly divergent non-sarbecovirus sequences had ≤27.1% amino acid identity to SARS-CoV-2 in ORF3a.

For the remaining 394 sequences, the ORF3c amino acid sequences were determined based on the conserved initiation site. In one (MG772933) ORF3c began with a GUG instead of an AUG codon. ORF3c was truncated in eight sequences. Five of these (EU371560, EU371561, EU371562, EU371563 and EU371564; ORF3c truncated to MLLLQVLFMLQQ) form a discrete subclade of SARS-CoV-1 viruses, but the sequences lack metadata and their provenance is unclear; nonetheless their ORF3a proteins have 99.3% amino acid identity to other SARS-CoV-1 isolates that have intact ORF3c. Another one (FJ882963; ORF3c truncated to MLLLQVLFML) also has an ORF3a protein with 99.3% amino acid identity to other SARS-CoV-1 isolates with intact ORF3c. Although these eight SARS-CoV-1 sequences were found to be ORF3c-defective, 184 other SARS-CoV-1 sequences had an ORF3c amino acid sequence that was identical to that of the SARS-CoV-1 reference sequence. In contrast, the remaining two sequences with truncated ORF3c (shown in Figure S1B) represent distinct sarbecovirus lineages.

ORF3c sequences with 100% amino acid identity to ORF3c of either the SARS-CoV-1 or SARS-CoV-2 NCBI reference sequences (NC_004718 and NC_045512, respectively) were discarded. Furthermore, four additional sequences (KF514407, AY463059, AY463060 and HG994853) with ≥99% amino acid identity to NC_004718 or NC_045512 in ORF3a were also discarded. In general, sequences with <99% identity in ORF3a were associated with non-human hosts and – where they had a different ORF3c amino acid sequence from NC_004718 and NC_045512 – were retained. Lab mutant recombinant sequences MT782114 and MT782115 were also removed. The remaining 191 ORF3c amino acid sequences represented 54 unique sequences (Figure S1A). ORF3c sequences were clustered with BLASTCLUST with a 90% identity threshold (-p T -L 0.95 -b T -S 90), resulting in seven clusters, and a representative sequence (SARS-CoV-1, SARS-CoV-2, or the most abundant unique sequence in the cluster) was chosen for Figure 1A.

### Analysis of SARS-CoV-2 ORF3c sequences from the GISAID database

The 13,427,526 available SARS-CoV-2 genome sequences were downloaded from epicov.org^77^, most recently on 10 Oct 2022. The findings of this study are based on sequence and metadata associated with 13,467,316 sequences available on GISAID via gisaid.org/EPI_SET_221101zc. All genome sequences and associated metadata in this dataset are published in GISAID’s EpiCoV database. To view the contributors of each individual sequence with details such as accession number, virus name, collection date, originating lab and submitting lab and the list of authors, visit https://doi.org/10.55876/gis8.221101zc.

To identify and extract ORF3c sequences, we searched for exact matches to any of seven 18-nt seed sequence queries spanning the region from the start of the ORF3a sgmRNA transcription regulatory sequence (TRS) to the nucleotide 5′-adjacent to the ORF3c AUG initiation codon, where each seed sequence had a 9-nt overlap with the previous seed sequence (viz. ACGAACTTATGGATTTGT, …, AGCAAGGTGAAATCAAGG). We then extracted the 200 nt downstream from the 3′-most matched seed, removed sequences with any “N”s (i.e. ambiguous nucleotides) in this region, trimmed the 5′ ends to the start of ORF3c, truncated the 3′ end at 126 nt downstream to encompass the 41 sense codons and the stop codon of ORF3c, leaving 12,906,225 sequences covering the ORF3c region.

These were translated and the number of occurrences of each unique 42-mer (amino acids + stop codons) were enumerated. Any sequence with fewer than 12,906 occurrences (i.e. 0.1% of total) were discarded, leaving 11 unique 42-mers. Five of these were variants of the PTC mutant where the different 42-mers give rise to a single translatable peptide MLLL, and therefore their occurrence counts (4,110,409, 21,375, 17,287, 15,798, and 14,876) were summed. The remaining seven variants are shown in Figure S6B. Collection date and country metadata were extracted from the sequence records and used to produce Figures 7D and 7E.

## FIGURE LEGENDS

**Supplemental Figure 1. Sarbecovirus ORF3c sequences. Related to Figure 1**.

(A) Unique ORF3c sequences. The number of times that each sequence was observed among the 191 available sarbecovirus ORF3c sequences is indicated at left and a representative accession number is shown at right. Host species from which viruses encoding the given ORF3c polypeptide have been sequenced are listed at far right. SARS-CoV-1 and SARS-CoV-2 sequences were excluded except for a single reference sequence for each.

(B) Two sequences (from two different studies) isolated from *Rhinolophus hipposideros* bats have a truncated or non-functional ORF3c that lacks the transmembrane region.

**Supplemental Figure 2. ORF3c mitochondrial distribution does not alter over time. Related to Figure 3**. Vero cells were transfected with a plasmid expressing ORF3c-HA. Cells were fixed at the indicated times and co-stained using antibodies against the HA epitope tag and either Tom20 (left panel) or Tom70 (right panel).

**Supplemental Figure 3. Interaction partners of ORF3c-HA were identified by SILAC LC-MS/MS**. Stable-isotope labelled Vero cells were transfected with a plasmid expressing ORF3c-HA or a HA-only control. Anti-HA immunoprecipitation was performed at 24 h post-transfection. Vertical lines indicate 0.5 log2 (i.e. 1.41-fold) enrichment or loss, and the horizontal dashed line represents a p-value of 0.05 (see Methods); q-values and FDR values of the six highlighted genes were also < 0.05. Grey markers indicate identified/quantified proteins. Blue markers indicate significantly enriched proteins, and the red marker indicates PGAM5.

**Supplemental Figure 4. ORF3c inhibits IFN-**β **signalling. Related to Figure 4**.

(A) ORF3c inhibits IFN-β and AP-1 reporter expression in a dose dependent manner. HEK293T cells were co-transfected with plasmids encoding IFN-β or AP-1-driven FLuc, as well as RLuc and an increasing dose of the ORF3c-HA plasmid. The cells were stimulated, and cell lysates were collected and analysed as described in Figure 4A.

(B) IFN-β and (C) AP-1 reporter gene assays as described above were performed with FLAG-tagged ORF3c, ORF3a, NSP6, NSP9, NSP12 or empty vector (EV). Statistical analysis (unpaired two sample t-test): ns = not significant, *P ≤ 0.05, **P ≤ 0.01, ***P ≤ 0.001, ****P ≤ 0.0001.

**Supplemental Figure 5. ORF3c co-localises with MAVS but not RIG-I, MDA5 or TRAF3 in SeV-infected cells. Related to Figure 5**.

(A) HEK293T cells (5 × 10^5^ cells) were seeded on coverslips pre-treated with 50 µg/mL poly-D-lysine. The next day, cells were transfected with 0.5 µg of plasmids expressing either FLAG-MAVS (upper panel), ORF3c-HA (second panel), both FLAG-MAVS and ORF3c-HA (third panel) or C6-HA and FLAG-MAVS (bottom panel). After 24 h incubation, cells were fixed and stained with mouse anti-HA, rabbit anti-FLAG and Mitotracker Red.

(B) HEK293T cells were transfected with 0.5 µg (total DNA) of plasmid expressing ORF3c-HA and FLAG-tagged MAVS, MDA5, RIG-I-CARD or TRAF3. Cells were fixed and stained as described in (A).

**Supplemental Figure 6. ORF3c absence does not alter mitochondrial morphology in infected cells. Related to Figure 7**.

(A) A549 +ACE2 +TMPRSS2 cells were infected with either early lineage (VIC1, B.1.1.7) or delta lineage (B.1.617.2-POR1, B.1.617.2-POR2) isolates and fixed at 24 h post infection. Cells were stained using antibodies against ORF3a and Tom20.

(B) Variants of ORF3c present in 0.1% or more of the 12,906,225 SARS-CoV-2 ORF3c sequences analysed. The number of occurrences is shown at left. Differences from the reference sequence (first sequence) are highlighted. See Figure 7D for variant abundances over time.

## SUPPLEMENTARY TABLES

**Supplemental Table 1**

MaxQuant proteinGroups output file for all quantified proteins from the Vero cell SILAC ORF3c-HA immunoprecipitation experiment after removing reverse database hits and contaminants.

**Supplemental Table 2**

Curated MaxQuant proteinGroups output file for the six quantified proteins that met >0.5 log2 (i.e. 1.41-fold) enrichment and p-value < 0.05 criteria for enrichment over HA-only control in the Vero cell ORF3c-HA SILAC immunoprecipitation experiment.

## REFERENCES

1. Finkel, Y., Mizrahi, O., Nachshon, A., Weingarten-Gabbay, S., Morgenstern, D., Yahalom-Ronen, Y., Tamir, H., Achdout, H., Stein, D., Israeli, O., et al. (2021). The coding capacity of SARS-CoV-2. Nature 589, 125–130. 10.1038/s41586-020-2739-1.

2. Jungreis, I., Sealfon, R., and Kellis, M. (2021). SARS-CoV-2 gene content and COVID-19 mutation impact by comparing 44 Sarbecovirus genomes. Nat Commun 12, 2642. 10.1038/s41467-021-22905-7.

3. Nelson, C.W., Ardern, Z., Goldberg, T.L., Meng, C., Kuo, C.H., Ludwig, C., Kolokotronis, S.O., and Wei, X. (2020). Dynamically evolving novel overlapping gene as a factor in the SARS-CoV-2 pandemic. Elife 9. 10.7554/eLife.59633.

4. Pavesi, A. (2020). New insights into the evolutionary features of viral overlapping genes by discriminant analysis. Virology 546, 51–66. 10.1016/j.virol.2020.03.007.

5. Konno, Y., Kimura, I., Uriu, K., Fukushi, M., Irie, T., Koyanagi, Y., Sauter, D., Gifford, R.J., Consortium, U.-C., Nakagawa, S., and Sato, K. (2020). SARS-CoV-2 ORF3b Is a Potent Interferon Antagonist Whose Activity Is Increased by a Naturally Occurring Elongation Variant. Cell Rep 32, 108185. 10.1016/j.celrep.2020.108185.

6. Gordon, D.E., Jang, G.M., Bouhaddou, M., Xu, J., Obernier, K., White, K.M., O’Meara, M.J., Rezelj, V.V., Guo, J.Z., Swaney, D.L., et al. (2020). A SARS-CoV-2 protein interaction map reveals targets for drug repurposing. Nature 583, 459–468. 10.1038/s41586-020-2286-9.

7. Michel, C.J., Mayer, C., Poch, O., and Thompson, J.D. (2020). Characterization of accessory genes in coronavirus genomes. Virol J 17, 131. 10.1186/s12985-020-01402-1.

8. Kim, D., Lee, J.Y., Yang, J.S., Kim, J.W., Kim, V.N., and Chang, H. (2020). The Architecture of SARS-CoV-2 Transcriptome. Cell 181, 914–921 e910. 10.1016/j.cell.2020.04.011.

9. Davidson, A.D., Williamson, M.K., Lewis, S., Shoemark, D., Carroll, M.W., Heesom, K.J., Zambon, M., Ellis, J., Lewis, P.A., Hiscox, J.A., and Matthews, D.A. (2020). Characterisation of the transcriptome and proteome of SARS-CoV-2 reveals a cell passage induced in-frame deletion of the furin-like cleavage site from the spike glycoprotein. Genome Med 12, 68. 10.1186/s13073-020-00763-0.

10. Firth, A.E. (2020). A putative new SARS-CoV protein, 3c, encoded in an ORF overlapping ORF3a. J Gen Virol 101, 1085–1089. 10.1099/jgv.0.001469.

11. Cagliani, R., Forni, D., Clerici, M., and Sironi, M. (2020). Coding potential and sequence conservation of SARS-CoV-2 and related animal viruses. Infect Genet Evol 83, 104353. 10.1016/j.meegid.2020.104353.

12. Jungreis, I., Nelson, C.W., Ardern, Z., Finkel, Y., Krogan, N.J., Sato, K., Ziebuhr, J., Stern-Ginossar, N., Pavesi, A., Firth, A.E., et al. (2021). Conflicting and ambiguous names of overlapping ORFs in the SARS-CoV-2 genome: A homology-based resolution. Virology 558, 145–151. 10.1016/j.virol.2021.02.013.

13. Delaune, D., Hul, V., Karlsson, E.A., Hassanin, A., Ou, T.P., Baidaliuk, A., Gambaro, F., Prot, M., Tu, V.T., Chea, S., et al. (2021). A novel SARS-CoV-2 related coronavirus in bats from Cambodia. Nat Commun 12, 6563. 10.1038/s41467-021-26809-4.

14. Zhou, H., Ji, J., Chen, X., Bi, Y., Li, J., Wang, Q., Hu, T., Song, H., Zhao, R., Chen, Y., et al. (2021). Identification of novel bat coronaviruses sheds light on the evolutionary origins of SARS-CoV-2 and related viruses. Cell 184, 4380–4391 e4314. 10.1016/j.cell.2021.06.008.

15. Wacharapluesadee, S., Tan, C.W., Maneeorn, P., Duengkae, P., Zhu, F., Joyjinda, Y., Kaewpom, T., Chia, W.N., Ampoot, W., Lim, B.L., et al. (2021). Evidence for SARS-CoV-2 related coronaviruses circulating in bats and pangolins in Southeast Asia. Nat Commun 12, 972. 10.1038/s41467-021-21240-1.

16. Murakami, S., Kitamura, T., Suzuki, J., Sato, R., Aoi, T., Fujii, M., Matsugo, H., Kamiki, H., Ishida, H., Takenaka-Uema, A., et al. (2020). Detection and Characterization of Bat Sarbecovirus Phylogenetically Related to SARS-CoV-2, Japan. Emerg Infect Dis 26, 3025–3029. 10.3201/eid2612.203386.

17. Lei, X., Dong, X., Ma, R., Wang, W., Xiao, X., Tian, Z., Wang, C., Wang, Y., Li, L., Ren, L., et al. (2020). Activation and evasion of type I interferon responses by SARS-CoV-2. Nat Commun 11, 3810. 10.1038/s41467-020-17665-9.

18. Lee, J.S., and Shin, E.C. (2020). The type I interferon response in COVID-19: implications for treatment. Nat Rev Immunol 20, 585–586. 10.1038/s41577-020-00429-3.

19. Takeuchi, O., and Akira, S. (2010). Pattern recognition receptors and inflammation. Cell 140, 805–820. 10.1016/j.cell.2010.01.022.

20. Cui, S., Eisenacher, K., Kirchhofer, A., Brzozka, K., Lammens, A., Lammens, K., Fujita, T., Conzelmann, K.K., Krug, A., and Hopfner, K.P. (2008). The C-terminal regulatory domain is the RNA 5’-triphosphate sensor of RIG-I. Mol Cell 29, 169–179. 10.1016/j.molcel.2007.10.032.

21. Pichlmair, A., Schulz, O., Tan, C.P., Rehwinkel, J., Kato, H., Takeuchi, O., Akira, S., Way, M., Schiavo, G., and Reis e Sousa, C. (2009). Activation of MDA5 requires higher-order RNA structures generated during virus infection. J Virol 83, 10761–10769. 10.1128/JVI.00770-09.

22. Zust, R., Cervantes-Barragan, L., Habjan, M., Maier, R., Neuman, B.W., Ziebuhr, J., Szretter, K.J., Baker, S.C., Barchet, W., Diamond, M.S., et al. (2011). Ribose 2’-O-methylation provides a molecular signature for the distinction of self and non-self mRNA dependent on the RNA sensor Mda5. Nat Immunol 12, 137–143. 10.1038/ni.1979.

23. Reikine, S., Nguyen, J.B., and Modis, Y. (2014). Pattern Recognition and Signaling Mechanisms of RIG-I and MDA5. Front Immunol 5, 342. 10.3389/fimmu.2014.00342.

24. Wathelet, M.G., Lin, C.H., Parekh, B.S., Ronco, L.V., Howley, P.M., and Maniatis, T. (1998). Virus infection induces the assembly of coordinately activated transcription factors on the IFN-beta enhancer in vivo. Mol Cell 1, 507–518. 10.1016/s1097-2765(00)80051-9.

25. Fu, Y.Z., Wang, S.Y., Zheng, Z.Q., Yi, H., Li, W.W., Xu, Z.S., and Wang, Y.Y. (2021). SARS-CoV-2 membrane glycoprotein M antagonizes the MAVS-mediated innate antiviral response. Cell Mol Immunol 18, 613–620. 10.1038/s41423-020-00571-x.

26. Jiang, H.W., Zhang, H.N., Meng, Q.F., Xie, J., Li, Y., Chen, H., Zheng, Y.X., Wang, X.N., Qi, H., Zhang, J., et al. (2020). SARS-CoV-2 Orf9b suppresses type I interferon responses by targeting TOM70. Cell Mol Immunol 17, 998–1000. 10.1038/s41423-020-0514-8.

27. Wu, J., Shi, Y., Pan, X., Wu, S., Hou, R., Zhang, Y., Zhong, T., Tang, H., Du, W., Wang, L., et al. (2021). SARS-CoV-2 ORF9b inhibits RIG-I-MAVS antiviral signaling by interrupting K63-linked ubiquitination of NEMO. Cell Rep 34, 108761. 10.1016/j.celrep.2021.108761.

28. Li, X., Hou, P., Ma, W., Wang, X., Wang, H., Yu, Z., Chang, H., Wang, T., Jin, S., Wang, X., et al. (2022). SARS-CoV-2 ORF10 suppresses the antiviral innate immune response by degrading MAVS through mitophagy. Cell Mol Immunol 19, 67–78. 10.1038/s41423-021-00807-4.

29. Schaecher, S.R., Mackenzie, J.M., and Pekosz, A. (2007). The ORF7b protein of severe acute respiratory syndrome coronavirus (SARS-CoV) is expressed in virus-infected cells and incorporated into SARS-CoV particles. J Virol 81, 718–731. 10.1128/JVI.01691-06.

30. Xu, K., Zheng, B.J., Zeng, R., Lu, W., Lin, Y.P., Xue, L., Li, L., Yang, L.L., Xu, C., Dai, J., et al. (2009). Severe acute respiratory syndrome coronavirus accessory protein 9b is a virion-associated protein. Virology 388, 279–285. 10.1016/j.virol.2009.03.032.

31. Firth, A.E., and Brierley, I. (2012). Non-canonical translation in RNA viruses. J Gen Virol 93, 1385–1409. 10.1099/vir.0.042499-0.

32. Kozak, M. (1986). Point mutations define a sequence flanking the AUG initiator codon that modulates translation by eukaryotic ribosomes. Cell 44, 283–292. 10.1016/0092-8674(86)90762-2.

33. Angelini, M.M., Akhlaghpour, M., Neuman, B.W., and Buchmeier, M.J. (2013). Severe acute respiratory syndrome coronavirus nonstructural proteins 3, 4, and 6 induce double-membrane vesicles. mBio 4. 10.1128/mBio.00524-13.

34. Santerre, M., Arjona, S.P., Allen, C.N., Shcherbik, N., and Sawaya, B.E. (2021). Why do SARS-CoV-2 NSPs rush to the ER? J Neurol 268, 2013–2022. 10.1007/s00415-020-10197-8.

35. Netland, J., Ferraro, D., Pewe, L., Olivares, H., Gallagher, T., and Perlman, S. (2007). Enhancement of murine coronavirus replication by severe acute respiratory syndrome coronavirus protein 6 requires the N-terminal hydrophobic region but not C-terminal sorting motifs. J Virol 81, 11520–11525. 10.1128/JVI.01308-07.

36. Frieman, M., Yount, B., Heise, M., Kopecky-Bromberg, S.A., Palese, P., and Baric, R.S. (2007). Severe acute respiratory syndrome coronavirus ORF6 antagonizes STAT1 function by sequestering nuclear import factors on the rough endoplasmic reticulum/Golgi membrane. J Virol 81, 9812–9824. 10.1128/JVI.01012-07.

37. Nelson, C.A., Pekosz, A., Lee, C.A., Diamond, M.S., and Fremont, D.H. (2005). Structure and intracellular targeting of the SARS-coronavirus Orf7a accessory protein. Structure 13, 75–85. 10.1016/j.str.2004.10.010.

38. Schaecher, S.R., Diamond, M.S., and Pekosz, A. (2008). The transmembrane domain of the severe acute respiratory syndrome coronavirus ORF7b protein is necessary and sufficient for its retention in the Golgi complex. J Virol 82, 9477–9491. 10.1128/JVI.00784-08.

39. Meier, C., Aricescu, A.R., Assenberg, R., Aplin, R.T., Gilbert, R.J., Grimes, J.M., and Stuart, D.I. (2006). The crystal structure of ORF-9b, a lipid binding protein from the SARS coronavirus. Structure 14, 1157–1165. 10.1016/j.str.2006.05.012.

40. Shi, C.S., Qi, H.Y., Boularan, C., Huang, N.N., Abu-Asab, M., Shelhamer, J.H., and Kehrl, J.H. (2014). SARS-coronavirus open reading frame-9b suppresses innate immunity by targeting mitochondria and the MAVS/TRAF3/TRAF6 signalosome. J Immunol 193, 3080–3089. 10.4049/jimmunol.1303196.

41. Ghosh, S., Dellibovi-Ragheb, T.A., Kerviel, A., Pak, E., Qiu, Q., Fisher, M., Takvorian, P.M., Bleck, C., Hsu, V.W., Fehr, A.R., et al. (2020). beta-Coronaviruses Use Lysosomes for Egress Instead of the Biosynthetic Secretory Pathway. Cell 183, 1520–1535 e1514. 10.1016/j.cell.2020.10.039.

42. Padhan, K., Tanwar, C., Hussain, A., Hui, P.Y., Lee, M.Y., Cheung, C.Y., Peiris, J.S.M., and Jameel, S. (2007). Severe acute respiratory syndrome coronavirus Orf3a protein interacts with caveolin. J Gen Virol 88, 3067–3077. 10.1099/vir.0.82856-0.

43. Tan, Y.J., Teng, E., Shen, S., Tan, T.H., Goh, P.Y., Fielding, B.C., Ooi, E.E., Tan, H.C., Lim, S.G., and Hong, W. (2004). A novel severe acute respiratory syndrome coronavirus protein, U274, is transported to the cell surface and undergoes endocytosis. J Virol 78, 6723–6734. 10.1128/JVI.78.13.6723-6734.2004.

44. Schoeman, D., and Fielding, B.C. (2019). Coronavirus envelope protein: current knowledge. Virol J 16, 69. 10.1186/s12985-019-1182-0.

45. Nieva, J.L., Madan, V., and Carrasco, L. (2012). Viroporins: structure and biological functions. Nat Rev Microbiol 10, 563–574. 10.1038/nrmicro2820.

46. Kemper, C., Habib, S.J., Engl, G., Heckmeyer, P., Dimmer, K.S., and Rapaport, D. (2008). Integration of tail-anchored proteins into the mitochondrial outer membrane does not require any known import components. J Cell Sci 121, 1990–1998. 10.1242/jcs.024034.

47. Chio, U.S., Cho, H., and Shan, S.O. (2017). Mechanisms of Tail-Anchored Membrane Protein Targeting and Insertion. Annu Rev Cell Dev Biol 33, 417–438. 10.1146/annurev-cellbio-100616-060839.

48. Knoops, K., Kikkert, M., Worm, S.H., Zevenhoven-Dobbe, J.C., van der Meer, Y., Koster, A.J., Mommaas, A.M., and Snijder, E.J. (2008). SARS-coronavirus replication is supported by a reticulovesicular network of modified endoplasmic reticulum. PLoS Biol 6, e226. 10.1371/journal.pbio.0060226.

49. Snijder, E.J., Limpens, R., de Wilde, A.H., de Jong, A.W.M., Zevenhoven-Dobbe, J.C., Maier, H.J., Faas, F., Koster, A.J., and Barcena, M. (2020). A unifying structural and functional model of the coronavirus replication organelle: Tracking down RNA synthesis. PLoS Biol 18, e3000715. 10.1371/journal.pbio.3000715.

50. Stertz, S., Reichelt, M., Spiegel, M., Kuri, T., Martinez-Sobrido, L., Garcia-Sastre, A., Weber, F., and Kochs, G. (2007). The intracellular sites of early replication and budding of SARS-coronavirus. Virology 361, 304–315. 10.1016/j.virol.2006.11.027.

51. Figueiredo Costa, B., Cassella, P., Colombo, S.F., and Borgese, N. (2018). Discrimination between the endoplasmic reticulum and mitochondria by spontaneously inserting tail-anchored proteins. Traffic 19, 182–197. 10.1111/tra.12550.

52. Costello, J.L., Castro, I.G., Camoes, F., Schrader, T.A., McNeall, D., Yang, J., Giannopoulou, E.A., Gomes, S., Pogenberg, V., Bonekamp, N.A., et al. (2017). Predicting the targeting of tail-anchored proteins to subcellular compartments in mammalian cells. J Cell Sci 130, 1675–1687. 10.1242/jcs.200204.

53. O’Keefe, S., Roboti, P., Duah, K.B., Zong, G., Schneider, H., Shi, W.Q., and High, S. (2021). Ipomoeassin-F inhibits the in vitro biogenesis of the SARS-CoV-2 spike protein and its host cell membrane receptor. J Cell Sci 134. 10.1242/jcs.257758.

54. O’Keefe, S., Zong, G., Duah, K.B., Andrews, L.E., Shi, W.Q., and High, S. (2021). An alternative pathway for membrane protein biogenesis at the endoplasmic reticulum. Commun Biol 4, 828. 10.1038/s42003-021-02363-z.

55. Wilson, R., Allen, A.J., Oliver, J., Brookman, J.L., High, S., and Bulleid, N.J. (1995). The translocation, folding, assembly and redox-dependent degradation of secretory and membrane proteins in semi-permeabilized mammalian cells. Biochem J 307 (Pt 3), 679–687. 10.1042/bj3070679.

56. Farkas, A., and Bohnsack, K.E. (2021). Capture and delivery of tail-anchored proteins to the endoplasmic reticulum. J Cell Biol 220. 10.1083/jcb.202105004.

57. Yu, Y.Q., Zielinska, M., Li, W., Bernkopf, D.B., Heilingloh, C.S., Neurath, M.F., and Becker, C. (2020). PGAM5-MAVS interaction regulates TBK1/ IRF3 dependent antiviral responses. Sci Rep 10, 8323. 10.1038/s41598-020-65155-1.

58. Yang, Z., Zheng, H., Li, H., Chen, Y., Hou, D., Fan, Q., Song, J., Guo, L., and Liu, L. (2021). The expression of IFN-beta is suppressed by the viral 3D polymerase via its impact on PGAM5 expression during enterovirus D68 infection. Virus Res 304, 198549. 10.1016/j.virusres.2021.198549.

59. Wang, Z., Jiang, H., Chen, S., Du, F., and Wang, X. (2012). The mitochondrial phosphatase PGAM5 functions at the convergence point of multiple necrotic death pathways. Cell 148, 228–243. 10.1016/j.cell.2011.11.030.

60. Lo, S.C., and Hannink, M. (2008). PGAM5 tethers a ternary complex containing Keap1 and Nrf2 to mitochondria. Exp Cell Res 314, 1789–1803. 10.1016/j.yexcr.2008.02.014.

61. Ma, K., Zhang, Z., Chang, R., Cheng, H., Mu, C., Zhao, T., Chen, L., Zhang, C., Luo, Q., Lin, J., et al. (2020). Dynamic PGAM5 multimers dephosphorylate BCL-xL or FUNDC1 to regulate mitochondrial and cellular fate. Cell Death Differ 27, 1036–1051. 10.1038/s41418-019-0396-4.

62. Chaikuad, A., Filippakopoulos, P., Marcsisin, S.R., Picaud, S., Schroder, M., Sekine, S., Ichijo, H., Engen, J.R., Takeda, K., and Knapp, S. (2017). Structures of PGAM5 Provide Insight into Active Site Plasticity and Multimeric Assembly. Structure 25, 1089–1099 e1083. 10.1016/j.str.2017.05.020.

63. Ruiz, K., Thaker, T.M., Agnew, C., Miller-Vedam, L., Trenker, R., Herrera, C., Ingaramo, M., Toso, D., Frost, A., and Jura, N. (2019). Functional role of PGAM5 multimeric assemblies and their polymerization into filaments. Nat Commun 10, 531. 10.1038/s41467-019-08393-w.

64. Seth, R.B., Sun, L., Ea, C.K., and Chen, Z.J. (2005). Identification and characterization of MAVS, a mitochondrial antiviral signaling protein that activates NF-kappaB and IRF 3. Cell 122, 669–682. 10.1016/j.cell.2005.08.012.

65. Hou, F., Sun, L., Zheng, H., Skaug, B., Jiang, Q.X., and Chen, Z.J. (2011). MAVS forms functional prion-like aggregates to activate and propagate antiviral innate immune response. Cell 146, 448–461. 10.1016/j.cell.2011.06.041.

66. Zhang, H., Zheng, H., Zhu, J., Dong, Q., Wang, J., Fan, H., Chen, Y., Zhang, X., Han, X., Li, Q., et al. (2021). Ubiquitin-Modified Proteome of SARS-CoV-2-Infected Host Cells Reveals Insights into Virus-Host Interaction and Pathogenesis. J Proteome Res 20, 2224–2239. 10.1021/acs.jproteome.0c00758.

67. Yuen, C.K., Lam, J.Y., Wong, W.M., Mak, L.F., Wang, X., Chu, H., Cai, J.P., Jin, D.Y., To, K.K., Chan, J.F., et al. (2020). SARS-CoV-2 nsp13, nsp14, nsp15 and orf6 function as potent interferon antagonists. Emerg Microbes Infect 9, 1418–1428. 10.1080/22221751.2020.1780953.

68. Li, J.Y., Liao, C.H., Wang, Q., Tan, Y.J., Luo, R., Qiu, Y., and Ge, X.Y. (2020). The ORF6, ORF8 and nucleocapsid proteins of SARS-CoV-2 inhibit type I interferon signaling pathway. Virus Res 286, 198074. 10.1016/j.virusres.2020.198074.

69. Xia, H., Cao, Z., Xie, X., Zhang, X., Chen, J.Y., Wang, H., Menachery, V.D., Rajsbaum, R., and Shi, P.Y. (2020). Evasion of Type I Interferon by SARS-CoV-2. Cell Rep 33, 108234. 10.1016/j.celrep.2020.108234.

70. Zheng, Y., Zhuang, M.W., Han, L., Zhang, J., Nan, M.L., Zhan, P., Kang, D., Liu, X., Gao, C., and Wang, P.H. (2020). Severe acute respiratory syndrome coronavirus 2 (SARS-CoV-2) membrane (M) protein inhibits type I and III interferon production by targeting RIG-I/MDA-5 signaling. Signal Transduct Target Ther 5, 299. 10.1038/s41392-020-00438-7.

71. Li, A., Zhao, K., Zhang, B., Hua, R., Fang, Y., Jiang, W., Zhang, J., Hui, L., Zheng, Y., Li, Y., et al. (2021). SARS-CoV-2 NSP12 Protein Is Not an Interferon-beta Antagonist. J Virol 95, e0074721. 10.1128/JVI.00747-21.

72. Clement, J.F., Bibeau-Poirier, A., Gravel, S.P., Grandvaux, N., Bonneil, E., Thibault, P., Meloche, S., and Servant, M.J. (2008). Phosphorylation of IRF-3 on Ser 339 generates a hyperactive form of IRF-3 through regulation of dimerization and CBP association. J Virol 82, 3984–3996. 10.1128/JVI.02526-07.

73. Pertel, T., Hausmann, S., Morger, D., Zuger, S., Guerra, J., Lascano, J., Reinhard, C., Santoni, F.A., Uchil, P.D., Chatel, L., et al. (2011). TRIM5 is an innate immune sensor for the retrovirus capsid lattice. Nature 472, 361–365. 10.1038/nature09976.

74. Li, X.D., Sun, L., Seth, R.B., Pineda, G., and Chen, Z.J. (2005). Hepatitis C virus protease NS3/4A cleaves mitochondrial antiviral signaling protein off the mitochondria to evade innate immunity. Proc Natl Acad Sci U S A 102, 17717–17722. 10.1073/pnas.0508531102.

75. Feng, H., Sander, A.L., Moreira-Soto, A., Yamane, D., Drexler, J.F., and Lemon, S.M. (2019). Hepatovirus 3ABC proteases and evolution of mitochondrial antiviral signaling protein (MAVS). J Hepatol 71, 25–34. 10.1016/j.jhep.2019.02.020.

76. Ning, X., Wang, Y., Jing, M., Sha, M., Lv, M., Gao, P., Zhang, R., Huang, X., Feng, J.M., and Jiang, Z. (2019). Apoptotic Caspases Suppress Type I Interferon Production via the Cleavage of cGAS, MAVS, and IRF3. Mol Cell 74, 19–31 e17. 10.1016/j.molcel.2019.02.013.

77. Elbe, S., and Buckland-Merrett, G. (2017). Data, disease and diplomacy: GISAID’s innovative contribution to global health. Glob Chall 1, 33–46. 10.1002/gch2.1018.

78. Sekine, S., Kanamaru, Y., Koike, M., Nishihara, A., Okada, M., Kinoshita, H., Kamiyama, M., Maruyama, J., Uchiyama, Y., Ishihara, N., et al. (2012). Rhomboid protease PARL mediates the mitochondrial membrane potential loss-induced cleavage of PGAM5. J Biol Chem 287, 34635–34645. 10.1074/jbc.M112.357509.

79. Yamaguchi, A., Ishikawa, H., Furuoka, M., Yokozeki, M., Matsuda, N., Tanimura, S., and Takeda, K. (2019). Cleaved PGAM5 is released from mitochondria depending on proteasome-mediated rupture of the outer mitochondrial membrane during mitophagy. J Biochem 165, 19–25. 10.1093/jb/mvy077.

80. Takeda, K., Komuro, Y., Hayakawa, T., Oguchi, H., Ishida, Y., Murakami, S., Noguchi, T., Kinoshita, H., Sekine, Y., Iemura, S., et al. (2009). Mitochondrial phosphoglycerate mutase 5 uses alternate catalytic activity as a protein serine/threonine phosphatase to activate ASK1. Proc Natl Acad Sci U S A 106, 12301–12305. 10.1073/pnas.0901823106.

81. Cheng, M., Lin, N., Dong, D., Ma, J., Su, J., and Sun, L. (2021). PGAM5: A crucial role in mitochondrial dynamics and programmed cell death. Eur J Cell Biol 100, 151144. 10.1016/j.ejcb.2020.151144.

82. Lin, H.Y., Lai, R.H., Lin, S.T., Lin, R.C., Wang, M.J., Lin, C.C., Lee, H.C., Wang, F.F., and Chen, J.Y. (2013). Suppressor of cytokine signaling 6 (SOCS6) promotes mitochondrial fission via regulating DRP1 translocation. Cell Death Differ 20, 139–153. 10.1038/cdd.2012.106.

83. He, G.W., Gunther, C., Kremer, A.E., Thonn, V., Amann, K., Poremba, C., Neurath, M.F., Wirtz, S., and Becker, C. (2017). PGAM5-mediated programmed necrosis of hepatocytes drives acute liver injury. Gut 66, 716–723. 10.1136/gutjnl-2015-311247.

84. Moriwaki, K., Farias Luz, N., Balaji, S., De Rosa, M.J., O’Donnell, C.L., Gough, P.J., Bertin, J., Welsh, R.M., and Chan, F.K. (2016). The Mitochondrial Phosphatase PGAM5 Is Dispensable for Necroptosis but Promotes Inflammasome Activation in Macrophages. J Immunol 196, 407–415. 10.4049/jimmunol.1501662.

85. Ishida, Y., Sekine, Y., Oguchi, H., Chihara, T., Miura, M., Ichijo, H., and Takeda, K. (2012). Prevention of apoptosis by mitochondrial phosphatase PGAM5 in the mushroom body is crucial for heat shock resistance in Drosophila melanogaster. PLoS One 7, e30265. 10.1371/journal.pone.0030265.

86. Xu, W., Jing, L., Wang, Q., Lin, C.C., Chen, X., Diao, J., Liu, Y., and Sun, X. (2015). Bax-PGAM5L-Drp1 complex is required for intrinsic apoptosis execution. Oncotarget 6, 30017–30034. 10.18632/oncotarget.5013.

87. Wu, H., Xue, D., Chen, G., Han, Z., Huang, L., Zhu, C., Wang, X., Jin, H., Wang, J., Zhu, Y., et al. (2014). The BCL2L1 and PGAM5 axis defines hypoxia-induced receptor-mediated mitophagy. Autophagy 10, 1712–1725. 10.4161/auto.29568.

88. Zhuang, M., Guan, S., Wang, H., Burlingame, A.L., and Wells, J.A. (2013). Substrates of IAP ubiquitin ligases identified with a designed orthogonal E3 ligase, the NEDDylator. Mol Cell 49, 273–282. 10.1016/j.molcel.2012.10.022.

89. Chen, G., Han, Z., Feng, D., Chen, Y., Chen, L., Wu, H., Huang, L., Zhou, C., Cai, X., Fu, C., et al. (2014). A regulatory signaling loop comprising the PGAM5 phosphatase and CK2 controls receptor-mediated mitophagy. Mol Cell 54, 362–377. 10.1016/j.molcel.2014.02.034.

90. Park, Y.S., Choi, S.E., and Koh, H.C. (2018). PGAM5 regulates PINK1/Parkin-mediated mitophagy via DRP1 in CCCP-induced mitochondrial dysfunction. Toxicol Lett 284, 120–128. 10.1016/j.toxlet.2017.12.004.

91. Lu, W., Karuppagounder, S.S., Springer, D.A., Allen, M.D., Zheng, L., Chao, B., Zhang, Y., Dawson, V.L., Dawson, T.M., and Lenardo, M. (2014). Genetic deficiency of the mitochondrial protein PGAM5 causes a Parkinson’s-like movement disorder. Nat Commun 5, 4930. 10.1038/ncomms5930.

92. Ganzleben, I., He, G.W., Gunther, C., Prigge, E.S., Richter, K., Rieker, R.J., Mougiakakos, D., Neurath, M.F., and Becker, C. (2019). PGAM5 is a key driver of mitochondrial dysfunction in experimental lung fibrosis. Cell Mol Life Sci 76, 4783–4794. 10.1007/s00018-019-03133-1.

93. Padhan, K., Minakshi, R., Towheed, M.A.B., and Jameel, S. (2008). Severe acute respiratory syndrome coronavirus 3a protein activates the mitochondrial death pathway through p38 MAP kinase activation. J Gen Virol 89, 1960–1969. 10.1099/vir.0.83665-0.

94. Ren, Y., Shu, T., Wu, D., Mu, J., Wang, C., Huang, M., Han, Y., Zhang, X.Y., Zhou, W., Qiu, Y., and Zhou, X. (2020). The ORF3a protein of SARS-CoV-2 induces apoptosis in cells. Cell Mol Immunol 17, 881–883. 10.1038/s41423-020-0485-9.

95. Bernkopf, D.B., Jalal, K., Bruckner, M., Knaup, K.X., Gentzel, M., Schambony, A., and Behrens, J. (2018). Pgam5 released from damaged mitochondria induces mitochondrial biogenesis via Wnt signaling. J Cell Biol 217, 1383–1394. 10.1083/jcb.201708191.

96. Mozzi, A., Oldani, M., Forcella, M.E., Vantaggiato, C., Cappelletti, G., Pontremoli, C., Valenti, F., Forni, D., Biasin, M., Sironi, M., et al. (2022). SARS-CoV-2 ORF3c impairs mitochondrial respiratory metabolism, oxidative stress and autophagic flow. bioRxiv, 2022.2010.2004.510754. 10.1101/2022.10.04.510754.

97. Kopecky-Bromberg, S.A., Martinez-Sobrido, L., Frieman, M., Baric, R.A., and Palese, P. (2007). Severe acute respiratory syndrome coronavirus open reading frame (ORF) 3b, ORF 6, and nucleocapsid proteins function as interferon antagonists. J Virol 81, 548–557. 10.1128/JVI.01782-06.

98. Miorin, L., Kehrer, T., Sanchez-Aparicio, M.T., Zhang, K., Cohen, P., Patel, R.S., Cupic, A., Makio, T., Mei, M., Moreno, E., et al. (2020). SARS-CoV-2 Orf6 hijacks Nup98 to block STAT nuclear import and antagonize interferon signaling. Proc Natl Acad Sci U S A 117, 28344–28354. 10.1073/pnas.2016650117.

99. Oh, S.J., and Shin, O.S. (2021). SARS-CoV-2 Nucleocapsid Protein Targets RIG-I-Like Receptor Pathways to Inhibit the Induction of Interferon Response. Cells 10. 10.3390/cells10030530.

100. Chen, K., Xiao, F., Hu, D., Ge, W., Tian, M., Wang, W., Pan, P., Wu, K., and Wu, J. (2020). SARS-CoV-2 Nucleocapsid Protein Interacts with RIG-I and Represses RIG-Mediated IFN-beta Production. Viruses 13. 10.3390/v13010047.

101. Menachery, V.D., Yount, B.L., Jr., Josset, L., Gralinski, L.E., Scobey, T., Agnihothram, S., Katze, M.G., and Baric, R.S. (2014). Attenuation and restoration of severe acute respiratory syndrome coronavirus mutant lacking 2’-o-methyltransferase activity. J Virol 88, 4251–4264. 10.1128/JVI.03571-13.

102. Hsu, J.C., Laurent-Rolle, M., Pawlak, J.B., Wilen, C.B., and Cresswell, P. (2021). Translational shutdown and evasion of the innate immune response by SARS-CoV-2 NSP14 protein. Proc Natl Acad Sci U S A 118. 10.1073/pnas.2101161118.

103. Kumar, A., Ishida, R., Strilets, T., Cole, J., Lopez-Orozco, J., Fayad, N., Felix-Lopez, A., Elaish, M., Evseev, D., Magor, K.E., et al. (2021). SARS-CoV-2 Nonstructural Protein 1 Inhibits the Interferon Response by Causing Depletion of Key Host Signaling Factors. J Virol 95, e0026621. 10.1128/JVI.00266-21.

104. Thorne, L.G., Bouhaddou, M., Reuschl, A.K., Zuliani-Alvarez, L., Polacco, B., Pelin, A., Batra, J., Whelan, M.V.X., Hosmillo, M., Fossati, A., et al. (2022). Evolution of enhanced innate immune evasion by SARS-CoV-2. Nature 602, 487–495. 10.1038/s41586-021-04352-y.

105. Gordon, D.E., Hiatt, J., Bouhaddou, M., Rezelj, V.V., Ulferts, S., Braberg, H., Jureka, A.S., Obernier, K., Guo, J.Z., Batra, J., et al. (2020). Comparative host-coronavirus protein interaction networks reveal pan-viral disease mechanisms. Science 370. 10.1126/science.abe9403.

106. Liu, X.Y., Wei, B., Shi, H.X., Shan, Y.F., and Wang, C. (2010). Tom70 mediates activation of interferon regulatory factor 3 on mitochondria. Cell Res 20, 994–1011. 10.1038/cr.2010.103.

107. Wei, B., Cui, Y., Huang, Y., Liu, H., Li, L., Li, M., Ruan, K.C., Zhou, Q., and Wang, C. (2015). Tom70 mediates Sendai virus-induced apoptosis on mitochondria. J Virol 89, 3804–3818. 10.1128/JVI.02959-14.

108. Edmonson, A.M., Mayfield, D.K., Vervoort, V., DuPont, B.R., and Argyropoulos, G. (2002). Characterization of a human import component of the mitochondrial outer membrane, TOMM70A. Cell Commun Adhes 9, 15–27. 10.1080/15419060212186.

109. Crozier, T., Greenwood, E., Williamson, J., Guo, W., Porter, L., Gabaev, I., Teixeira-Silva, A., Grice, G., Wickenhagen, A., Stanton, R., et al. (2022). Quantitative proteomic analysis of SARS-CoV-2 infection of primary human airway ciliated cells and lung epithelial cells demonstrates the effectiveness of SARS-CoV-2 innate immune evasion [version 1; peer review: awaiting peer review]. Wellcome Open Research 7. 10.12688/wellcomeopenres.17946.1.

110. Bojkova, D., Klann, K., Koch, B., Widera, M., Krause, D., Ciesek, S., Cinatl, J., and Munch, C. (2020). Proteomics of SARS-CoV-2-infected host cells reveals therapy targets. Nature 583, 469–472. 10.1038/s41586-020-2332-7.

111. Perrin-Cocon, L., Diaz, O., Jacquemin, C., Barthel, V., Ogire, E., Ramiere, C., Andre, P., Lotteau, V., and Vidalain, P.O. (2020). The current landscape of coronavirus-host protein-protein interactions. J Transl Med 18, 319. 10.1186/s12967-020-02480-z.

112. Meyer, B., Chiaravalli, J., Gellenoncourt, S., Brownridge, P., Bryne, D.P., Daly, L.A., Grauslys, A., Walter, M., Agou, F., Chakrabarti, L.A., et al. (2021). Characterising proteolysis during SARS-CoV-2 infection identifies viral cleavage sites and cellular targets with therapeutic potential. Nat Commun 12, 5553. 10.1038/s41467-021-25796-w.

113. Osada, N., Kohara, A., Yamaji, T., Hirayama, N., Kasai, F., Sekizuka, T., Kuroda, M., and Hanada, K. (2014). The genome landscape of the african green monkey kidney-derived vero cell line. DNA Res 21, 673–683. 10.1093/dnares/dsu029.

114. Yount, B., Roberts, R.S., Sims, A.C., Deming, D., Frieman, M.B., Sparks, J., Denison, M.R., Davis, N., and Baric, R.S. (2005). Severe acute respiratory syndrome coronavirus group-specific open reading frames encode nonessential functions for replication in cell cultures and mice. J Virol 79, 14909–14922. 10.1128/JVI.79.23.14909-14922.2005.

115. Castano-Rodriguez, C., Honrubia, J.M., Gutierrez-Alvarez, J., DeDiego, M.L., Nieto-Torres, J.L., Jimenez-Guardeno, J.M., Regla-Nava, J.A., Fernandez-Delgado, R., Verdia-Baguena, C., Queralt-Martin, M., et al. (2018). Role of Severe Acute Respiratory Syndrome Coronavirus Viroporins E, 3a, and 8a in Replication and Pathogenesis. mBio 9. 10.1128/mBio.02325-17.

116. Freundt, E.C., Yu, L., Goldsmith, C.S., Welsh, S., Cheng, A., Yount, B., Liu, W., Frieman, M.B., Buchholz, U.J., Screaton, G.R., et al. (2010). The open reading frame 3a protein of severe acute respiratory syndrome-associated coronavirus promotes membrane rearrangement and cell death. J Virol 84, 1097–1109. 10.1128/JVI.01662-09.

117. Rihn, S.J., Merits, A., Bakshi, S., Turnbull, M.L., Wickenhagen, A., Alexander, A.J.T., Baillie, C., Brennan, B., Brown, F., Brunker, K., et al. (2021). A plasmid DNA-launched SARS-CoV-2 reverse genetics system and coronavirus toolkit for COVID-19 research. PLoS Biol 19, e3001091. 10.1371/journal.pbio.3001091.

118. Caly, L., Druce, J., Roberts, J., Bond, K., Tran, T., Kostecki, R., Yoga, Y., Naughton, W., Taiaroa, G., Seemann, T., et al. (2020). Isolation and rapid sharing of the 2019 novel coronavirus (SARS-CoV-2) from the first patient diagnosed with COVID-19 in Australia. Med J Aust 212, 459–462. 10.5694/mja2.50569.

119. Lulla, V., Lulla, A., Wernike, K., Aebischer, A., Beer, M., and Roy, P. (2016). Assembly of Replication-Incompetent African Horse Sickness Virus Particles: Rational Design of Vaccines for All Serotypes. J Virol 90, 7405–7414. 10.1128/JVI.00548-16.

120. Lulla, V., Dinan, A.M., Hosmillo, M., Chaudhry, Y., Sherry, L., Irigoyen, N., Nayak, K.M., Stonehouse, N.J., Zilbauer, M., Goodfellow, I., and Firth, A.E. (2019). An upstream protein-coding region in enteroviruses modulates virus infection in gut epithelial cells. Nat Microbiol 4, 280–292. 10.1038/s41564-018-0297-1.

121. Unterholzner, L., Sumner, R.P., Baran, M., Ren, H., Mansur, D.S., Bourke, N.M., Randow, F., Smith, G.L., and Bowie, A.G. (2011). Vaccinia virus protein C6 is a virulence factor that binds TBK-1 adaptor proteins and inhibits activation of IRF3 and IRF7. PLoS Pathog 7, e1002247. 10.1371/journal.ppat.1002247.

122. Zhang, J., Cruz-Cosme, R., Zhuang, M.W., Liu, D., Liu, Y., Teng, S., Wang, P.H., and Tang, Q. (2020). A systemic and molecular study of subcellular localization of SARS-CoV-2 proteins. Signal Transduct Target Ther 5, 269. 10.1038/s41392-020-00372-8.

123. Soday, L., Lu, Y., Albarnaz, J.D., Davies, C.T.R., Antrobus, R., Smith, G.L., and Weekes, M.P. (2019). Quantitative Temporal Proteomic Analysis of Vaccinia Virus Infection Reveals Regulation of Histone Deacetylases by an Interferon Antagonist. Cell Rep 27, 1920–1933 e1927. 10.1016/j.celrep.2019.04.042.

124. Lulla, V., and Firth, A.E. (2020). A hidden gene in astroviruses encodes a viroporin. Nat Commun 11, 4070. 10.1038/s41467-020-17906-x.

125. McKenna, M., Simmonds, R.E., and High, S. (2016). Mechanistic insights into the inhibition of Sec61-dependent co- and post-translational translocation by mycolactone. J Cell Sci 129, 1404–1415. 10.1242/jcs.182352.

126. Roboti, P., Sato, K., and Lowe, M. (2015). The golgin GMAP-210 is required for efficient membrane trafficking in the early secretory pathway. J Cell Sci 128, 1595–1606. 10.1242/jcs.166710.

127. Hughes, C.S., Moggridge, S., Muller, T., Sorensen, P.H., Morin, G.B., and Krijgsveld, J. (2019). Single-pot, solid-phase-enhanced sample preparation for proteomics experiments. Nat Protoc 14, 68–85. 10.1038/s41596-018-0082-x.

128. Tyanova, S., Temu, T., and Cox, J. (2016). The MaxQuant computational platform for mass spectrometry-based shotgun proteomics. Nat Protoc 11, 2301–2319. 10.1038/nprot.2016.136.

129. Storey, J.D. (2002). A Direct Approach to False Discovery Rates. Journal of the Royal Statistical Society. Series B (Statistical Methodology) 64, 479–498.

130. Altschul, S.F., Gish, W., Miller, W., Myers, E.W., and Lipman, D.J. (1990). Basic local alignment search tool. J Mol Biol 215, 403–410. 10.1016/S0022-2836(05)80360-2.

131. Edgar, R.C. (2004). MUSCLE: multiple sequence alignment with high accuracy and high throughput. Nucleic Acids Res 32, 1792–1797. 10.1093/nar/gkh340.

132. Rice, P., Longden, I., and Bleasby, A. (2000). EMBOSS: the European Molecular Biology Open Software Suite. Trends Genet 16, 276–277. 10.1016/s0168-9525(00)02024-2.

133. Kall, L., Krogh, A., and Sonnhammer, E.L. (2007). Advantages of combined transmembrane topology and signal peptide prediction--the Phobius web server. Nucleic Acids Res 35, W429–432. 10.1093/nar/gkm256.

134. Wang, S., Li, W., Liu, S., and Xu, J. (2016). RaptorX-Property: a web server for protein structure property prediction. Nucleic Acids Res 44, W430–435. 10.1093/nar/gkw306.

