## Supplemental Figure 1 for "The SARS-CoV-2 protein ORF3c is a mitochondrial modulator of innate immunity"

**(A)**

| | | | |
|---|---|---|---|
| 14 | MLLLQVLFMLLQRYRYKPHSLSDGLLLVLHCFLFSRALP | OL674078 | *Rhinolophus stheno, R. malayanus, R. affinis* |
| 1 | MILLQILFALQQKYRYKPHSLSDGLLLVLHFLLFFRALPKS | LC663959 | *Rhinolophus cornutus* |
| 1 | MILLQILFALQQKYRYKPHSLSDGLLLVLHFLLFFKALPKL | LC556375 | *Rhinolophus cornutus* |
| 2 | MILLQILFALQQKYRYKPHSLSDGLLLVLHFLLFFKALPKS | LC663793 | *Rhinolophus cornutus* |
| 1 | VLLLQILFALLQQYRYKPHSLSDGLLLALQFLLFFKALQK | MG772933 | *Rhinolophus pusillus* |
| 1 | MLLLQILFALLQQYRYKPHPLSDGLLLALQFLLFFKALQK | OK017805 | *Rhinolophus pusillus* |
| 5 | MLLLQILFALLQQYRYKPHSLSDGLLLALQFLLFFKALQK | MW703458 | *Rhinolophus pusillus*, *R. blythi,* |
| 11 | MLLLQVLFMLLQRYHYKPHSLSDGLLLALHFLLFFKALLK | DQ022305 | *Rhinolophus sp.* |
| 1 | MLLLQVLFMLLQRYHYKPHSLSDGLLLALHFLLFFKALPQ | OK017792 | *Rhinolophus affinis* |
| 1 | MLLLQVLFMLLQRYHYRPHSLSDGLLLALHFLLFFKALPK | OK017838 | *Rhinolophus sinicus* |
| 1 | MLLLQILFALLQRYRYKPHSLSDGLLLALHFLLFFKALPRS | MN996532 | *Rhinolophus affinis* |
| 1 | MLLLQILFALLQRYRYKPHSLSDGLLLALHSLLFFKALPKS | OK287355 | *Rhinolophus blythi* |
| 1 | MLLLQILFALLQRYRYKPHSLSDGLLLALHFLLFFKALPK | MT121216 | *Manis javanica* |
| 4 | MLLLQILFALLQRYRYKPHSLSDGLLLALHFLLFFKALPKS | MZ937003 | *Rhinolophus malayanus*, *R. pusillus*, *R. marshalli* |
| 1 | MLLLQILFALLQRYRYKPHSLSDGLLLALHFLLFFTALPK | MT040333 | *Manis javanica* |
| 5 | MLLLQILFALLQRYRYKPHSLSDGLLLALHFLLFSRALPKL | MW251308 | *Rhinolophus acuminatus* |
| 1 | MLLLQILFALPQRYRYKPHSLSDGLLLALHFLLFFRALPKS | MZ937000 | *Rhinolophus malayanus* |
| 1 | MLLLQILFALLQRYRYKPHSLSDGLLLALHFLLFFRALPKS | NC_045512 | *Homo sapiens* (SARS-CoV-2) |
| 1 | MLLLQILFALLQRYRYKPHSLSDGLLLALHFLLFFRALPKL | MZ081381 | *Rhinolophus pusillus* |
| 6 | MLLLQILFALLQRYRYKPHSLSDGLLLALHFLLFFRALPK | MT040335 | *Manis javanica* |
| 1 | MFLLQVLFMLLQRYRYKPHSLSDGLLLALHFLLFFKALPN | MZ081382 | *Rhinolophus stheno* |
| 1 | MLLLQVLFMLQQRYRYRPHSPSDGLLLALHFLLFFKALPK | MZ328294 | *Rhinolophus sinicus* |
| 1 | MLLLQILFMLQQRYRYKPHSRSDGLLLALHFLLFFKALPK | OK017860 | *Rhinolophus siamensis* |
| 1 | MLLLQVLFMLLQRYRCKPHSPLDGLLLALHFLLFFKALPK | OK017812 | *Rhinolophus sinicus* |
| 1 | MLLLQVLFMLLQRYRYKPHSLLDGLLLALHFLLFFRALPK | KY417143 | *Rhinolophus sinicus* |
| 1 | MLLLQVLFMLLQRYRYKPHSRLDGLLLALHFLLFFKALPK | OK017859 | *Rhinolophus siamensis* |
| 1 | MLLLQVLFMLLQRYRYKPHSLLDGLLLALHFLLFFKALPK | MZ081377 | *Rhinolophus malayanus* |
| 1 | MLLLQVLFMLLQRYRYKPHSPLDGLLLALHFLLFFKALPK | OK017843 | *Rhinolophus sinicus* |
| 15 | MLLLQVLFMLLQRYRCKPHSPSDGLLLALHFLLFFKALPK | MW681002 | *Rhinolophus sinicus*, *R. affinis* |
| 1 | MLLLQVLFMLLQQYRYKPHSPSDGLLLALHFLLFFKALPK | FJ588690 | *Rhinolophus sinicus* |
| 1 | MLLLQVLFMLLQRYPYKPHSPSDGLLLALHFLLFFKALPK | OK017823 | *Rhinolophus sinicus* |
| 8 | MLLLQVLFMLLQRYRYKPHSPSDGLLLALHFLLFFKALPK | OK017801 | *Rhinolophus sinicus*, *R. affinis* |
| 3 | MLLLQVLFMLQQRYRYRPHSLSDGLLLALHFLLFFKALPR | DQ071615 | *Rhinolophus sinicus* |
| 1 | MLLLQVLFMLQQRYRYKPHSLSDGLLLALHFLLFFKALPR | OK017830 | *Rhinolophus sinicus* |
| 1 | MLLLQVLFMLQQRYRYKPHSHSDGLLLALHFLLFFKALPK | JX993987 | *Rhinolophus pusillus* |
| 6 | MLLLQVLFMLQQRYRYKPHSLSDGLLLALHFLLFFKALPK | DQ412043 | *Rhinolophus sinicus*, *R. macrotis* |
| 1 | MLLLQVLFMLLQRYRYKPHSLSDGLLLALHFLLFFRALPK | KY417148 | *Rhinolophus sinicus* |
| 1 | MLLLQVLFMLQQRYRYKPHSFSDGLLLALHFLLFFRALPK | AY304492 | |
| 15 | MLLMQVLFVLQQRYRYKPHSLSDGLLLALHFLLFFRALPK | AY545917 | *Paguma larvata*, *Melogale moschata* |
| 2 | MLLLQVLFVLQQRYRYKPHSLSDGLLLALHFLLFFRALPK | AY545916 | *Paguma larvata* |
| 8 | MLLLQVLFMLQQRYRYRPHSLSDGLLLALHFLLFFRALPK | MK211376 | *Rhinolophus sinicus*, *R. affinis* |
| 1 | MLLLQVLFMLQQRYRYKPHSLSDGLLLALHFLLFFRALPK | NC_004718 | *Homo sapiens* (SARS-CoV) |
| 2 | MLLLQVLFMLQQRYHYKPHSLSDGLLLALHFLLFFRALPK | KY417151 | *Rhinolophus sinicus* |
| 7 | MLLLQVLFMLQQRYHYRPHSLSDGLLLALHFLLFFRALPK | OK017858 | *Rhinolophus sinicus* |
| 1 | MLFLQVLFMLLQQYRYKPHSLSDGLLLALHFLLFFKALPR | ON378802 | *Rhinolophus ferrumequinum* |
| 1 | MLFLQVLFMLLQRYRYKPHSLSDGLLLALHFLLFFKALPR | KY938558 | *Rhinolophus ferrumequinum* |
| 1 | MLLLQVLFMLLQRYRYKPHSLSDGLLLALHFFLFFKALPK | KJ473815 | *Rhinolophus sinicus* |
| 21 | MLLLQVLFMLLQRYRYKPHSLSDGLLLALHFLLFFKALPK | KY417149 | *R. sinicus*, *R. affinis*, *R. ferrumequinum*, *Aselliscus stoliczkanus* |
| 1 | MLFLQVLFMLLQRYRYKPHSLSDGLLLVLHFLLFFKALPK | JX993988 | *Chaerephon plicata* |
| 20 | MLFLQVLFMLLQRYRYKPHSLSDGLLLALHFLLFFKALPK | KP886808 | *Rhinolophus ferrumequinum*, *R. sinicus* |
| 1 | MFLLQVLFILQRPSLYRPLYLSDGLLLALHFLLFFKALRN | NC_014470 | *Rhinolophus blasii* |
| 1 | MFLLQILFMLQLPSLYKPLYLSDGLLLALHCWLFFKTLQK | MZ190137 | *Rhinolophus ferrumequinum* |
| 1 | MLFLQILFMLRLPSLYKPLCLSDGLLLALHCLLFFKMLQK | KY352407 | *Rhinolophus sp.* |
| 2 | MLFLQILFMLRLPSLYKPLCLSDGLLLALHCLLFFKTLQK | MT726045 | *Rhinolophus sp.* |

```
  ::::*:** *        .*     *****.*:   **    *
```

**(B)**

| | | | |
|---|---|---|---|
| 1 | MLLPLVLSKLQPPYLYRPLCLLVGSL | MW719567 | *Rhinolophus hipposideros* |
| 1 | MYHLLVQYMPKPTFHYRHHCRLAGLL | MZ190138 | *Rhinolophus hipposideros* |
