## Supplementary figures and images for "The SARS-CoV-2 protein ORF3c is a mitochondrial modulator of innate immunity"

### Supplemental Figure 2

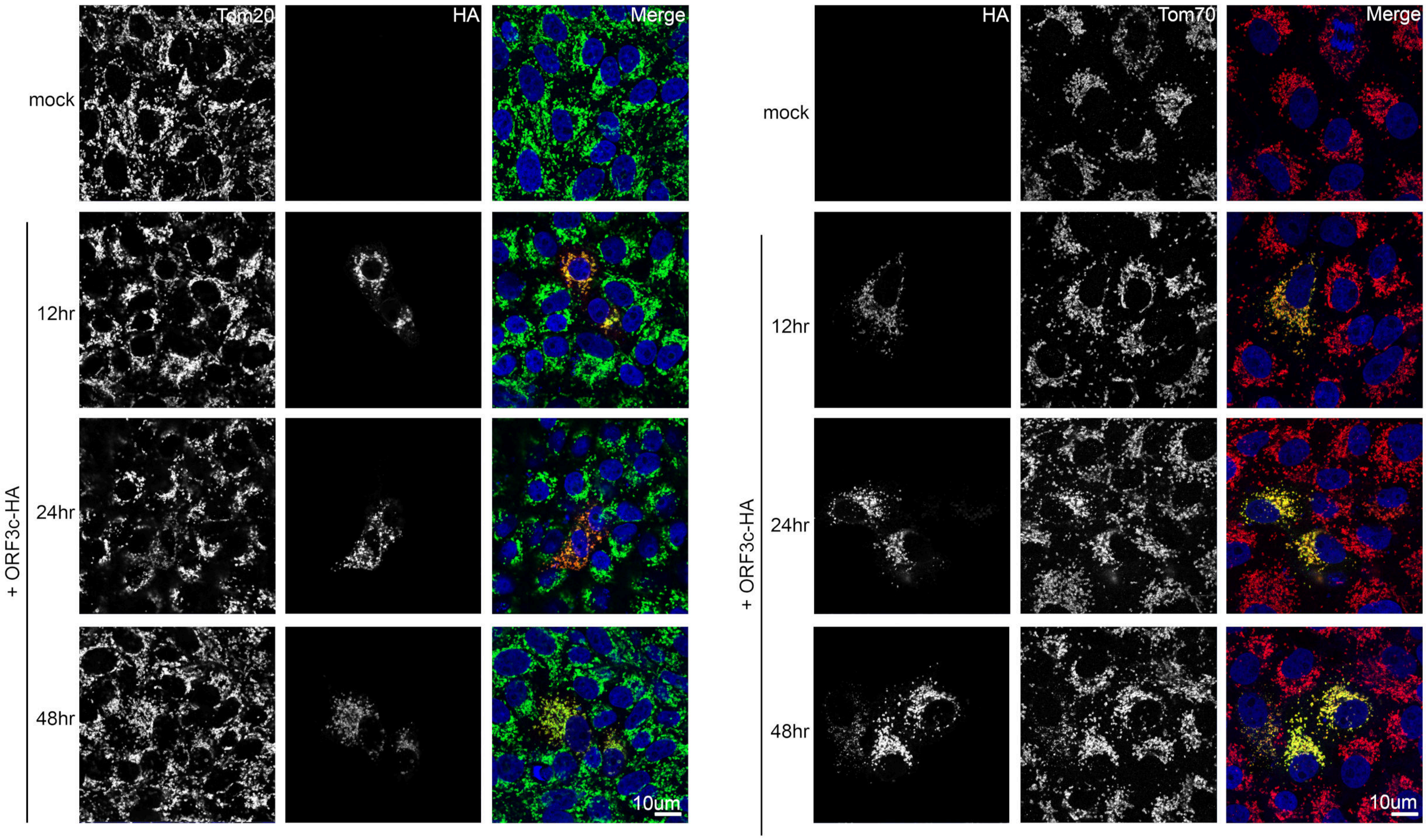

### Supplemental Figure 3

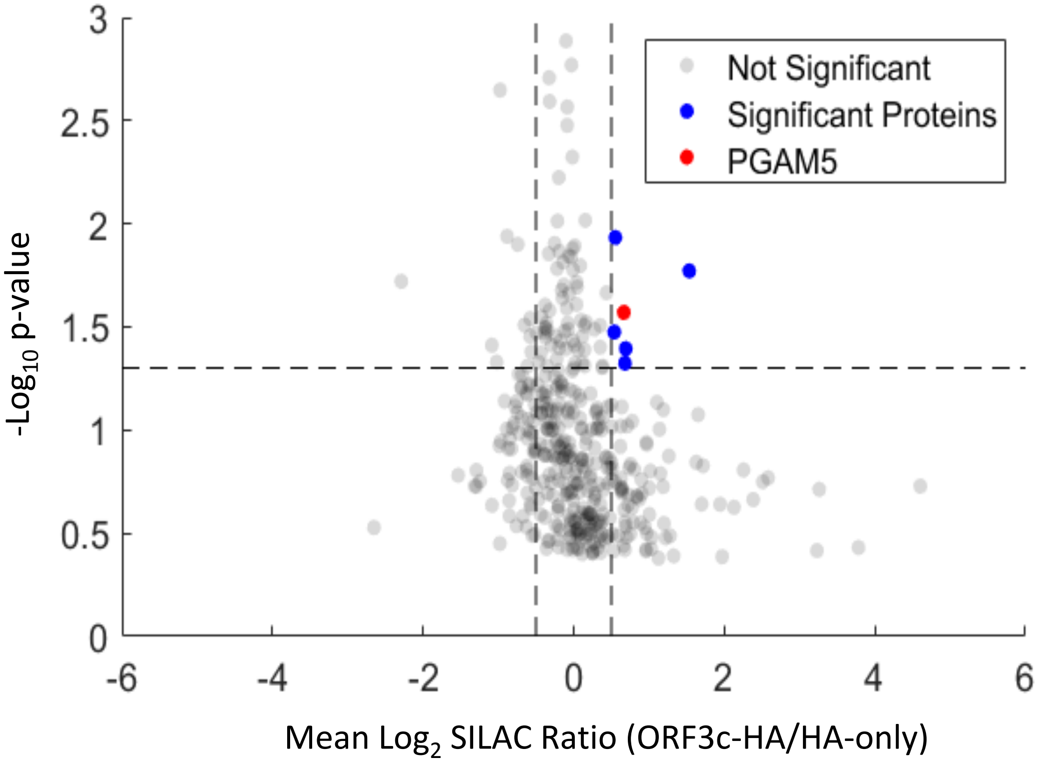

### Supplemental Figure 4

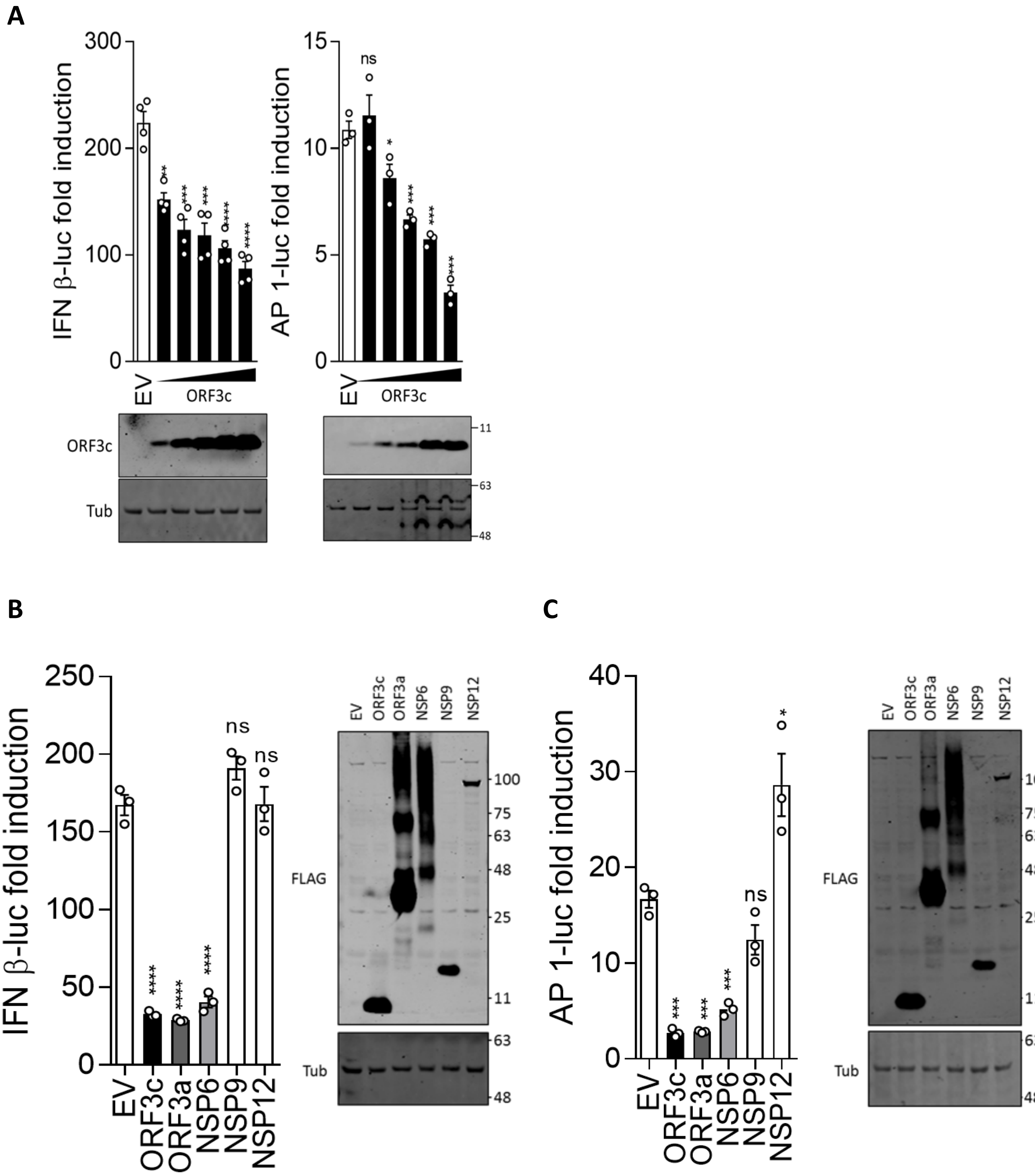

### Supplemental Figure 5

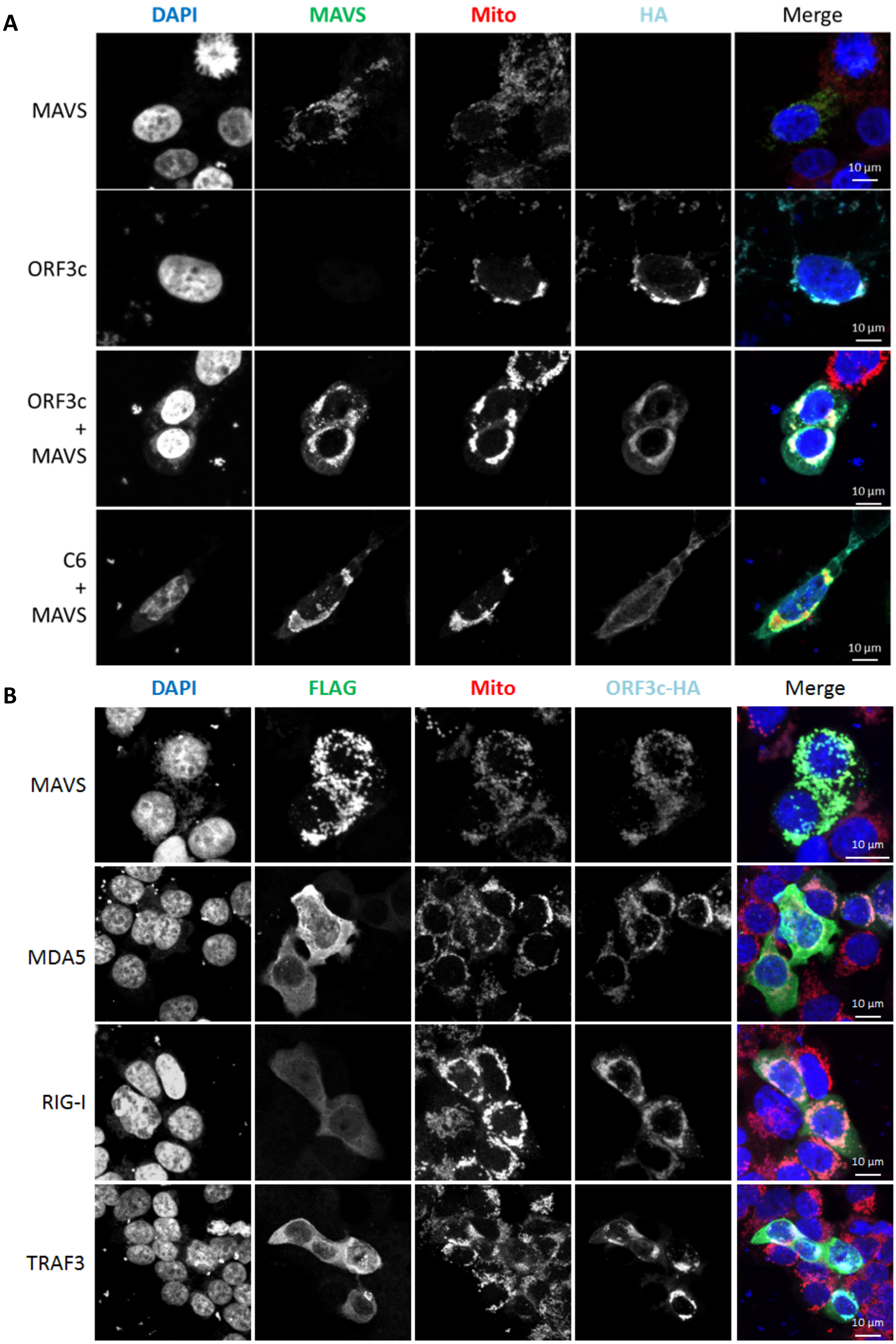
