## Supplemental Figure 6 for "The SARS-CoV-2 protein ORF3c is a mitochondrial modulator of innate immunity"

**A**

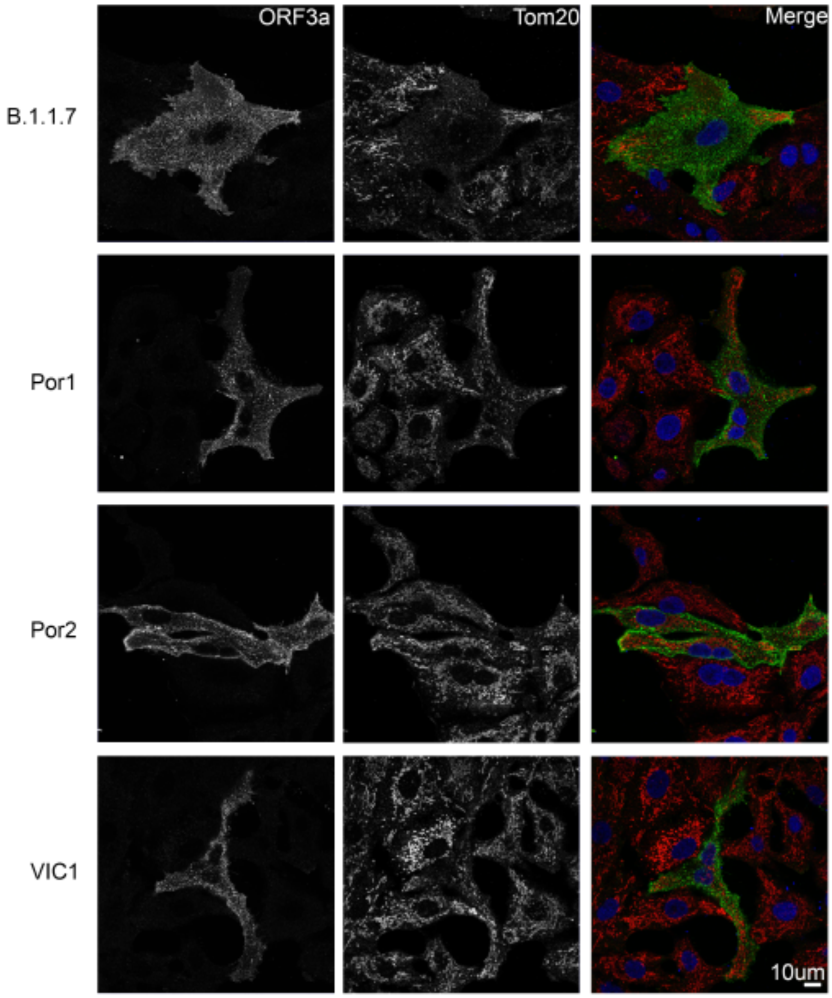


**B**

| | | |
|---|---|---|
| 7685950 | MLLLQILFALLQRYRYKPHSLSDGLLLALHFLLFFRALPKS | WT |
| 4179745 | MLLL | PTC |
| 481382 | MLLLQILFALLQRYRYKPHSLSDGLLLALHFLLFFIALPKS | R36I |
| 57355 | MLLLQILFALLQRYRYKPHSFSDGLLLALHFLLFFIALPKS | L21F/R36I |
| 34046 | MLLLQILFALLQRYRYKPHSLLDGLLLALHFLLFFRALPKS | S22L |
| 30815 | MLLLYILFALLQRYRYKPHSLSDGLLLALHFLLFFRALPKS | Q5Y |
| 15368 | MLLLQILFALLQRYRYEPHSLSDGLLLALHFLLFFRALPKS | K17E |
